# Targeting acetyl-CoA metabolism attenuates the formation of fear memories through reduced activity-dependent histone acetylation

**DOI:** 10.1101/2022.05.22.492937

**Authors:** Desi C Alexander, Tanya Corman, Mariel Mendoza, Andrew Glass, Tal Belity, Riane R Campbell, Joseph Han, Ashley A Keiser, Jeffrey Winkler, Marcelo A Wood, Thomas Kim, Benjamin A Garcia, Hagit Cohen, Philipp Mews, Gabor Egervari, Shelley L Berger

## Abstract

Histone acetylation is a key component in the consolidation of long-term fear memories. Epigenetic enzymes involved in histone acetylation, including histone acetyltransferases and deacetylases, have been put forward as potential pharmacological targets in the treatment of pathological fear memories, such as those that underlie post-traumatic stress disorder (PTSD). However, these enzymes typically play a ubiquitous role in gene regulation, which precludes the clinical use of systemic manipulations. Recently, we have found that a nuclear-localized metabolic enzyme, Acetyl-coA synthetase 2 (Acss2), modulates histone acetylation during learning and memory. Loss of Acss2 is well-tolerated in mice, with no impact on general health or baseline behavior. Here, we show that an Acss2 null mouse model shows reduced acquisition of long-term fear memories in assays of contextual and cued fear conditioning. We find that loss of Acss2 leads to consolidation-specific reductions in both histone acetylation and the expression of critical learning and memory-related genes in the dorsal hippocampus. Further, we show that systemic administration of blood-brain-barrier (BBB)-permeable Acss2 inhibitors during the consolidation window reduces fear memory formation in mice and rats, and also reduces anxiety in a predator-scent-stress (PSS) paradigm. Our findings suggest that Acss2 plays a critical role in the formation of fear memories, and represents a potential pharmacological target in the treatment of PTSD.

## INTRODUCTION

The ability to form robust, persistent memories in response to danger is essential for survival in all animals. However, the strength of these fear memories can come at a cost of long-term stress and associated disease. At its most severe, fear memory in humans can become pathological, resulting in post-traumatic stress disorder (PTSD). PTSD develops following exposure to traumatic events such as interpersonal violence, life-threatening accidents, or military combat when memories of a traumatic event are chronically and inappropriately reactivated, resulting in intrusive retrieval of traumatic memories, insomnia, and irritability^1^. Despite the overwhelming burden that PTSD places on affected individuals, their communities, and the health system, current treatment options are limited^2^. In fact, approximately 50% of patients fail to respond to trauma-based psychotherapy. While symptom improvement is experienced by half of the patients, two-thirds of those treated still meet the criteria for a PTSD diagnosis with adverse symptoms such as intrusive memories and emotional withdrawal^3,4^. Due to this, discovery of novel pharmacological targets for PTSD is a pressing public health concern^5^.

Histone acetylation is a major regulator of many types of long-term memory, including fear memories^6^. The formation of long-term memories (LTM) involves the physical remodeling of synapses between firing cells, requiring robust transcription of plasticity-dependent genes^7^. The initial wave of gene expression involves a set of genes called the immediate early genes (IEGs), which are upregulated within 30 minutes of neuronal stimulation and which initiate downstream gene expression^8^. The transcription of IEGs is facilitated by the deposition of histone acetylation at their promoters, and the subsequent removal of these activating marks results in downregulation of IEG transcripts^9^. Histone acetylation is balanced by the activity of histone acetyltransferases (HATs), such as CREB-binding protein (CBP), and histone deacetylases (HDACs), and this balance is of critical importance in the brain^9^.

Because of the key role histone acetylation plays in the formation of LTM, epigenetic proteins have become attractive targets for therapeutic memory manipulation. For example, Class I HDAC inhibitors are being actively pursued as enhancers of memory in mouse models of Alzheimer’s disease^10^. Inhibitors of CBP and/or its homologue p300, which reduce LTM in the dorsal hippocampus (dHPC) and basolateral amygdala (BLA), have been proposed as therapeutic agents for the treatment of PTSD. However, CBP is not an ideal target as null mice are embryonic lethal^11^. In humans, loss of a single allele of CBP or its paralogue p300 results in Rubinstein-Taybi syndrome, a condition characterized by severe intellectual disability^12,13^. The substrate for histone acetylation is the central metabolite acetyl-coA. Although synthesized in the mitochondria as part of central metabolism, acetyl-coA is produced throughout the cell, where it serves as a key substrate for the biosynthesis of lipids, ketones, and the neurotransmitter acetylcholine^14^, as well as for acetylation of proteins. Several acetyl-coA-producing enzymes translocate into the nucleus, where they aid in gene expression by producing a pool of acetyl-coA for direct use by histone acetyltransferases^15,16^. In the yeast *Saccharomyces cerevisiae*, this pool is mediated by the acetyl-coA synthetase ACS2, which converts acetate into acetyl-coA. Loss of ACS2 results in global deacetylation of histone tails, and subsequent downregulation of most transcription^17^. In contrast to yeast, most mammalian systems have moved away from acetate as a bioenergetic substrate, relying instead on the TCA intermediate citrate. In mammalian cells, citrate is harvested from the TCA cycle and converted to acetyl-coA through the enzyme ATP citrate lyase (ACLY). ACLY thus plays a critical role in maintaining histone acetylation in mammals, particularly in highly mitotic (and glycolytic) cancer cells^18^. Other metabolic enzymes also support histone acetylation under certain conditions. For example the pyruvate dehydrogenase complex (PDC) contributes to histone acetylation during early embryonic development, and the mammalian homologue of ACS2, acetyl-coA synthetase 2 (ACSS2) is required to maintain histone acetylation during metabolic stress, such as glucose starvation^19^. In contrast to PDC or ACLY, which rely on products of glucose metabolism (pyruvate and citrate, respectively), ACSS2 can recycle acetate released by histone deacetylase reactions. This capability for acetate reprocessing implies that ACSS2 may be of particular importance in post-mitotic cells like neurons, which do not dilute their nuclear metabolic pools through cell division.

We previously demonstrated that ACSS2 is critical for histone acetylation in the adult mouse brain, particularly during the establishment of long-term spatial memories^20,21^. Stereotactic knockdown of ACSS2 in the dorsal hippocampus (dHPC) of adult mice substantially impairs the formation of long-term spatial memories, consistent with reduced activity-dependent transcription in the dHPC. However, whether ACSS2 is required for highly durable forms of LTM, such as fear memory, is unknown. Moreover, whether ACSS2 represents a robust pharmacological target to ameliorate diseases of persistent traumatic memories, such as PTSD, remains a pressing question.

In this study, we examined the contribution of ACSS2 to formation of fear memory in rodent models. We utilized two approaches: a null ACSS2 mouse model and pharmacological inhibition of ACSS2 using small-molecule inhibitors. Our results support a crucial role for ACSS2 in fear memory via regulation of histone acetylation and transcription of key immediate early genes. Overall, our findings underscore a profound impact of a metabolic protein, ACSS2, on the epigenome and transcriptome in the learning brain and establish ACSS2 as a promising target in treating PTSD.

## RESULTS

### Generation of a constitutive ACSS2^KO^ mouse

To examine the role of ACSS2 in learning and memory, we used CRISPR/Cas9 to engineer mice with a deletion of exons 3-7 of the *Acss2* locus (*Acss2*Δ^3-7^). We designed guide RNAs (gRNAs) to target the Cas9 enzyme to intronic regions upstream of exon 3 and downstream of exon 7. The resulting deletion introduces multiple premature stop codons, with the first located at the new junction between exon 2 and exon 8 (Fig. 1a).

**Figure 1:**
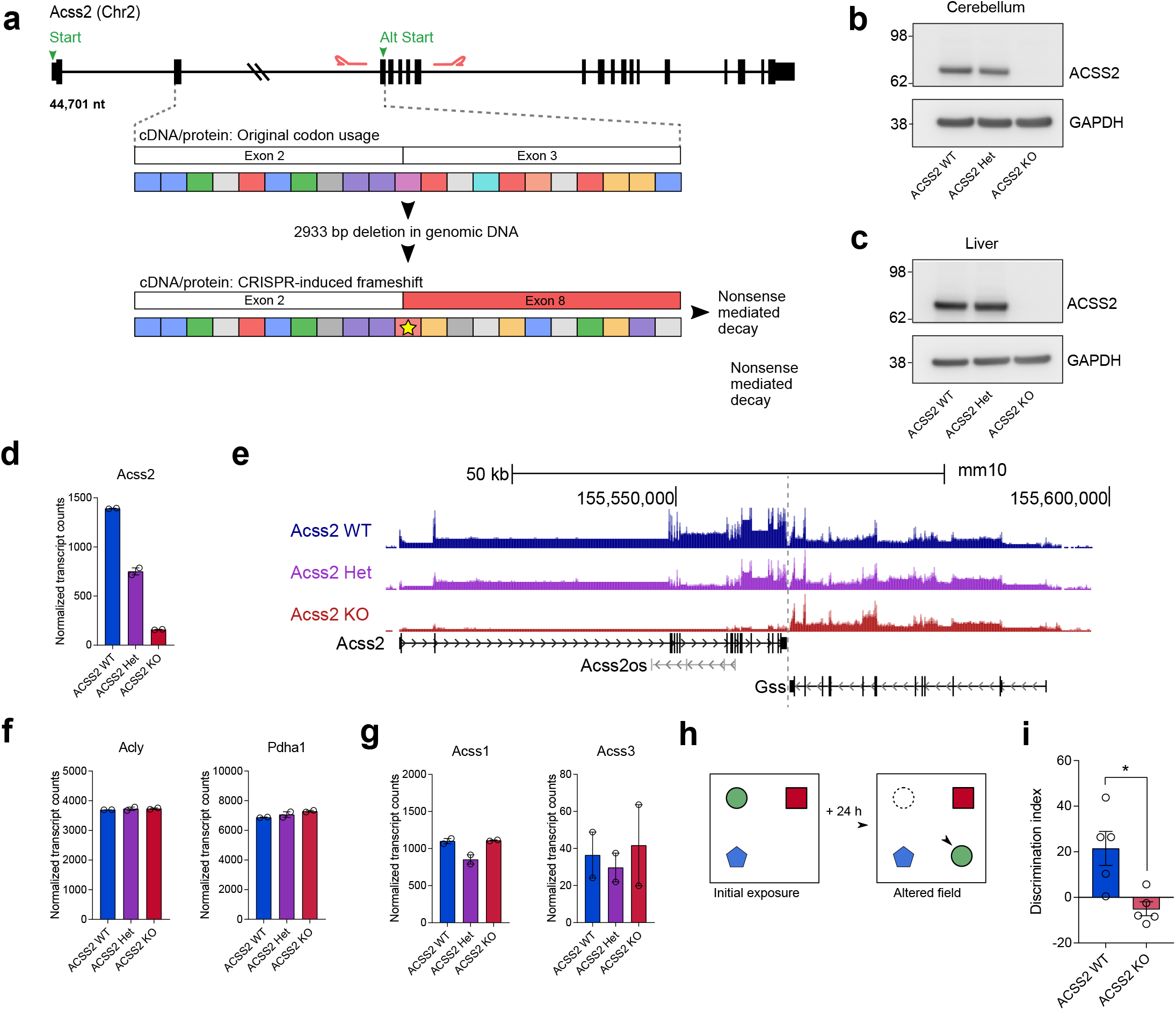
Generation of a constitutive Acss2KO mouse. **a)** Diagram showing CRISPR strategy for ACSS2 knockout in mice. Guide RNAs (gRNAs) shown as red hairpins; stop codons as yellow stars. **b)** Representative western blot of ACSS2 expression in brain (cerebellum), showing lysates from age-matched WT, Acss2het, and Acss2KO mice **c)** Representative western blot of ACSS2 expression in liver. **d)** Acss2 mRNA levels shown as normalized transcript counts mapped to Acss2 exons (RNA-seq, STAR, normalized with Deseq2). **e)** UCSC genome browser showing changing levels of Acss2 transcript (on left), and unchanged neighboring gene (Gss7) on right. **f)** Bar charts showing mRNA expression levels of Acly and PDC subunits (Pdha1, Pdhb, Pdhx, Dld, Dlat). **g)** Bar charts of Acss1 and Acss2 mRNA levels shown as normalized transcript counts. **h)** Object location memory (OLM) assay. **i)** Bar graph showing OLM discrimination score for WT and Acss2KO mice (WT: n=5; KO: n=5, p = 0.011, unpaired t-test).

We injected these guides along with Cas9 mRNA into fertilized eggs harvested from C57Bl6/J mice (Jackson Laboratories). To remove potential mutations derived from off-target Cas9 activity, we crossed founder *Acss2*Δ^3-7^ mice back to the founding C57BL6/J strain for three generations. A previously generated mouse homozygous null KO of ACSS2 was viable^22,23^; however, this strain was unavailable, and was a mixed C57Bl6/129 background, which reduces suitability for behavioral research

Mice harboring biallelic copies of *Acss2*Δ^3-7^ (hereafter called Acss2^KO^) exhibited complete loss of ACSS2 protein across all tissues, including the cerebellum (Fig. 1b) and liver (Fig. 1c). *Acss2* mRNA was ablated across the entire *Acss2* locus, including upstream of the 5’ cut site (Figs. 1d, 1e), likely through nonsense-mediated decay^24^. Importantly, the Acss2^KO^ line is viable, phenotypically normal under standard housing conditions, shows no premature mortality, and breeds consistently with no reductions in litter size, consistent with the previous ACSS2 null mouse^22,23^.

Loss of one enzymatic source of acetyl-coA can result in compensation from other acetyl-coA-producing enzymes^18^. Acetyl-CoA, as a central metabolite, can be produced by a number of enzymes, including ATP citrate lyase (ACLY) in the cytosol and pyruvate dehydrogenase complex (PDC) in the mitochondria; as mentioned above, both enzymes can translocate into the nucleus under certain conditions^18,25,26^. Within RNA-seq datasets, we found no increased transcription of *Acly* or any PDC components (including *Pdha1*, which encodes the active site of the E1 subunit) in the dorsal hippocampus of the Acss2^KO^ mice (Fig. 1f, Supplemental Fig. 1a), nor upregulation of other acetyl-coA synthetase family members (*Acss1, Acss3*) (Fig.1g). These results show that loss of Acss2 is well-tolerated in the brain under baseline conditions, without obvious re-wiring of acetyl-CoA metabolism.

### Acss2^KO^ mice are not behaviorally impaired under baseline conditions

The ACSS2 KO mice exhibited no visible morphological or behavioral abnormalities. We assayed the mice for mild neurological impairments using a battery of assays to test locomotion, anxiety, and working memory. The open field assay simultaneously measures locomotion and anxiety in rodents by quantifying distance traveled and affinity towards the periphery of the testing arena (a behavior known as thigmotaxis (Supplementary Fig. 1a). Representative traces of both WT and Acss2^KO^ mice generated by ANY-maze during this interval were visually similar (Supplementary Fig. 1b). We found that Acss2^KO^ mice exhibited no difference in path length (Supplementary Fig. 1c), or thigmotaxis (Supplementary Fig. 1d) compared to wild-type controls, demonstrating normal locomotion and anxiety.

We then examined working memory, which involves cytoskeletal rearrangements, receptor trafficking, and post-translational modifications of existing proteins at the firing synapse, i.e. processes that are upstream of transcription-dependent long-term memory consolidation^27^. To test whether Acss2^KO^ mice have intact working memory, we used a y-maze test of spontaneous alternation, which relies on correct recall of previous paths when choosing a new maze arm to explore^28^ (Supplemental Fig. 1e). ACSS2^KO^ mice showed no deficit, engaging in spontaneous alternations at wild-type levels (Supplemental Fig. 1f).

Our previous studies showed that knockdown of Acss2 in the dorsal hippocampus leads to a deficit in long-term spatial memory, such as object location memory (OLM)^20^ and conditioned place preference (CPP)^21^. To test whether this deficit holds true in the Acss2^KO^ mouse, we again tested OLM performance (Fig. 1h). Consistent with our previous results^20,21^, Acss2^KO^ mice exhibited significantly reduced discrimination index compared to wild-type controls (Fig. 1i), thus indicating poor consolidation of spatial memories.

### Loss of ACSS2 impairs fear conditioning

We tested long-term memory using a fear conditioning (FC) paradigm which pairs an aversive stimulus with a contextual or auditory cue^29^. We subjected age-matched ACSS2^KO^ and wild-type mice to a standard FC paradigm (Fig. 2a), placing mice in an arena for 2 minutes of exploration, then presenting an auditory cue coincident with a mild foot shock (1 mA, 2s); the mice were returned to the arena 24 hours after training to test context and cue (auditory) recall.

**Figure 2:**
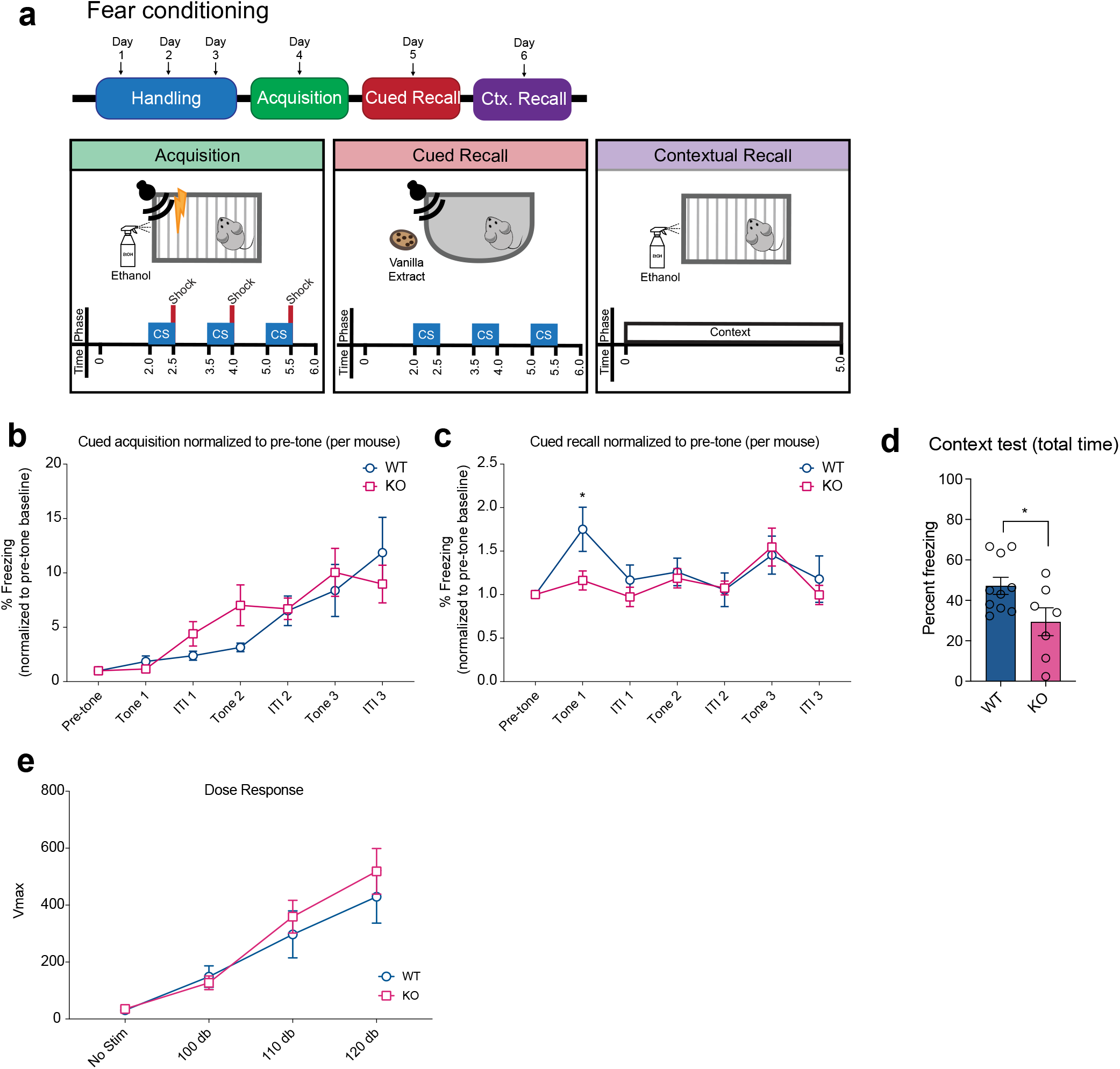
Loss of ACSS2 impairs fear conditioning. **a)** Fear conditioning schematic, showing training/acquisition (panel 1), cued/auditory recall (panel 2), and contextual recall (panel 3). **b)** Fear response during training/acquisition reflected as Fear response during training/acquisition reflected as fold change over freezing within the pre-tone interval. Periods binned by period – pre-tone, inter-tone interval (ITI) or tone. (p = n.s., 2-way ANOVA). **c)** Fear response during cued/auditory recall reflected as fold change over pre-tone interval. Periods binned by period – pre-tone, inter-tone interval (ITI) or tone. (2-way ANOVA, *post hoc* Fisher’s LSD *p=0.0208). **d)** Fear response during contextual recall reflected as percent time freezing, averaged over the entire recall period (5 min). (p = 0.0305, unpaired t-test). **e)** Input/output (dose response) curves generated by acoustic startle response assay (p=ns, 2-way ANOVA).

The previous results of ACSS2^KO^ mice maintaining working memory but having poor LTM (Supplemental Fig. 1 and Fig. 1), predicted that acquisition of fear memory may be normal in the ACSS2 KO mice, but that consolidation of memory may be defective. Analysis of freezing during training showed that Acss2^KO^ mice readily responded to FC in the short term, and exhibited increased freezing with every presentation of the tone-shock pair (Fig. 2b, Supplemental Fig. 3b). However, during recall sessions, Acss2^KO^ mice performed significantly worse than WT controls, where WT mice responded robustly to the onset of the first tone by increasing freezing 1.68-fold. Acss2^KO^ freezing was significantly reduced (Figs. 2c, Supplemental Fig. 3c, p = 0.02 of first tone). Ruling out auditory deficit in the Acss2^KO^ mouse as a potential confound, we found that the Acss2^KO^ mice performed at WT levels in an acoustic startle response assay (Fig. 2e).

Importantly, Acss2^KO^ mice also froze significantly less than WT mice during the contextual recall session (Fig. 2d, Supplementary Fig. 3d), which depends on the recall of spatial memories. Together, these results indicate that Acss2^KO^ mice are deficient in the consolidation of long-term fear memory.

### Acss2^KO^ mice exhibit impairment of histone acetylation relevant to long-term memory

Histone acetylation is required for the transcription of activity-dependent genes, and certain histone acetylation sites such as H4K8ac^30^, H4K5ac^31^, and H3K27ac^32^ are linked to activity-related gene expression in learning and memory models. We analyzed histone post-translational modifications (PTMs) during long-term memory formation in FC using “bottom-up” mass spectrometry which allows unbiased quantification of global histone PTMs^33^. We subjected an age-matched cohort of mice (4 months old; WT n=3, KO n=2) to contextual FC, using a single shock paradigm (1.5 mA, 2s) which generated long-term fear memories (Supplementary Figs. 3e, 3f, 3g) and provided a clearly defined starting point, allowing time-specific sample collection (Fig. 3a). To capture the first wave of activity-related histone acetylation, we sacrificed mice 30 minutes after acquisition (Fig. 3a). We analyzed the dorsal hippocampus because contextual FC relies on spatial memories formed there. Homecage-housed WT and Acss2^KO^ mice served as baseline controls (WT baseline: n=5, KO baseline n=3).

**Figure 3:**
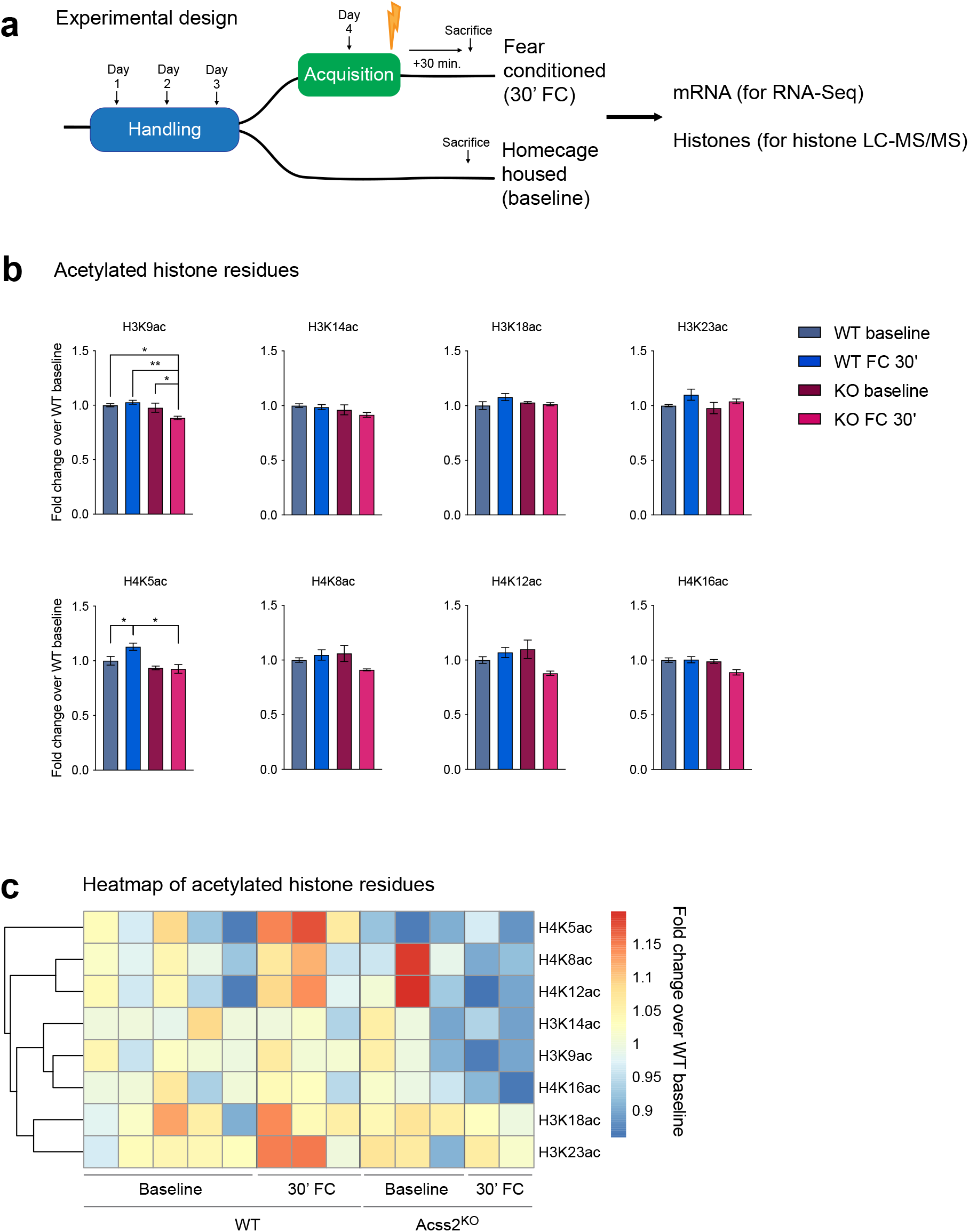
ACSS2 KO mice exhibit reduced histone acetylation in an activity-dependent context. **a)** Schematic of paired histone acetylation and RNA-seq study design. **b)** Bar graphs showing histone acetylation levels expressed as fold-change over WT baseline (abundance/average baseline abundance) ((*) p-value < 0.05, One-way ANOVA, *post hoc* Fisher’s LSD). **c)** Heatmap of all histone marks expressed as fold change over WT baseline.

In WT mice, we found significant increases in H4K5ac 30 minutes after FC (Fig. 3b, One-way ANOVA; p = 0.0238, Fisher’s LSD, p = 0.0304), consistent with previous reports of upregulation in the dorsal hippocampus (dHPC) after FC^31^. While there were no significant differences in histone acetylation between WT and ACSS2^KO^ mice at baseline (Fig. 3b), following FC, ACSS2^KO^ mice exhibited marked reductions in H3K9ac (One-way ANOVA, p = 0.0290), when compared to WT FC (Fisher’s LSD, p = 0.0055), and baseline KO (Fisher’s LSD, p = 0.0424) and H4K5ac (One-way ANOVA p = 0.0371) when compared to WT FC (Fisher’s LSD, p = 0.0102). While not achieving significance, other acetylation marks on the H4 tail (H4K8ac, H4K12ac, and H4K16ac) showed a similar downward trend. Unbiased clustering analysis indicated that not all histone acetylation sites were similarly affected by the loss of ACSS2; for example, H3K18ac and H3K23ac showed no fluctuations upon ACSS2 deletion (Fig. 3c).

### Acss2^KO^ mice exhibit transcriptional dysregulation in response to fear conditioning

Histone acetylation is required for the expression of activity-related genes in learning and memory^11,34^. Given the observed changes in global histone acetylation during FC, we next assayed transcriptional responses in the WT and Acss2^KO^ brain. RNA-seq was performed from slices of dHPC reserved from the same mice used to examine histone acetylation. A direct comparison between WT and ACSS2^KO^ mice after FC showed acute transcriptional alteration (Fig. 4a), with 2873 genes significantly differentially expressed between genotypes (DESeq2, padj < 0.05). The 1654 genes enriched in WT mice after FC featured gene ontology categories related to LTM formation, including learning, calcium ion transport, and transmembrane ion channels (Fig. 4c). Aside from Acss2 itself, the most significantly enriched genes in the WT dHPC after FC included proteins required for the remodeling of activated synapses, including the cytoskeletal protein Actinin Alpha 1 (Actn1)^35,36^, the ionotropic glutamate [NMDA] receptor subunit epsilon-2 (Grin2b)^37,38^, and nitric oxide synthetase 1 (Nos1), which generates nitric oxide at the post-synaptic density (PSD) for use as a second messenger, neurotransmitter, and vasodilator^39^. We also noted downregulation of the histone acetyltrasferases Crebbp (CBP), p300 and Kat2a (Gcn5) in the KO mouse; all three enzymes have been characterized in learning and memory models^34,40–42^ (Supplementary Fig. 4e).

**Figure 4:**
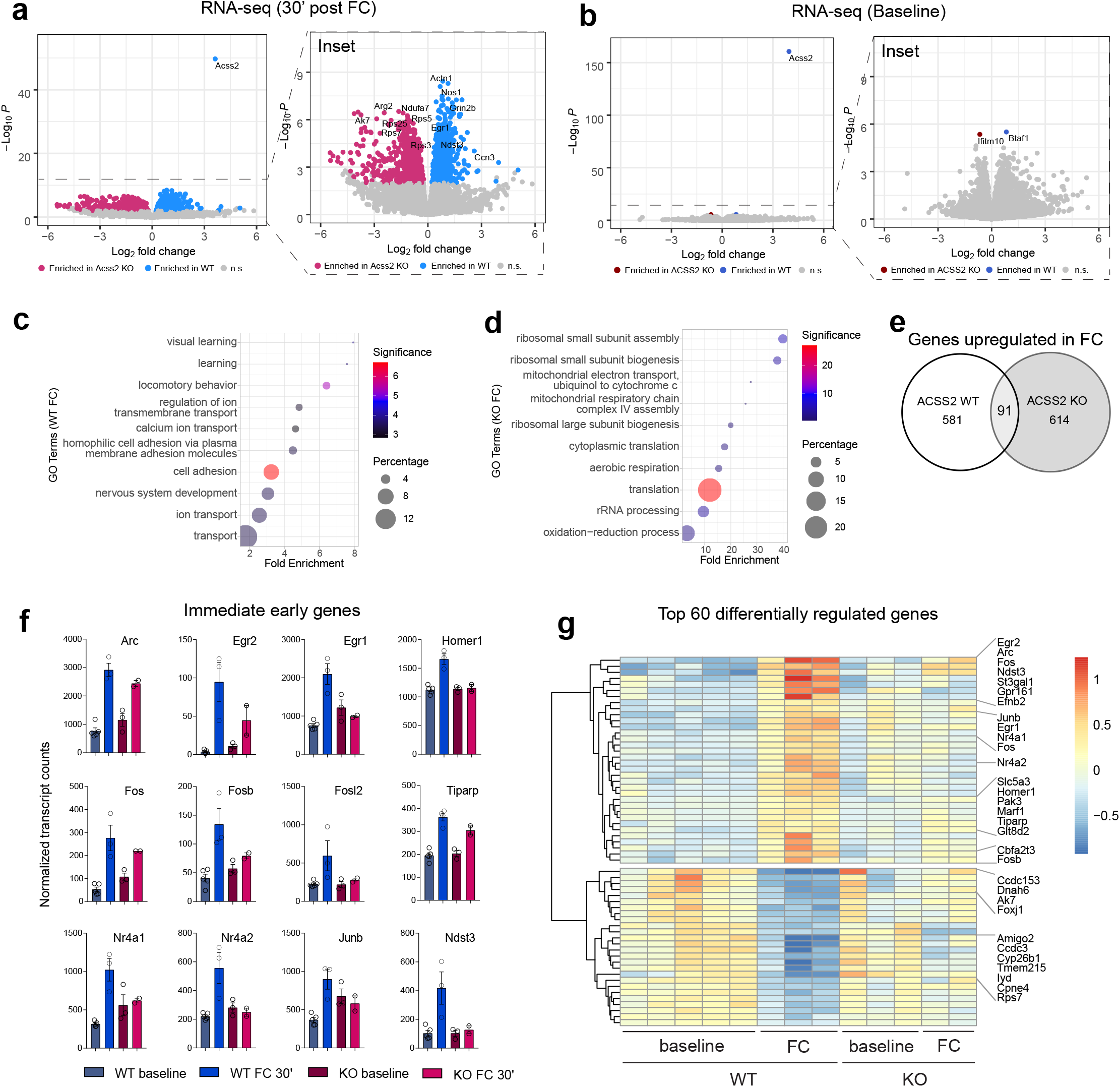
Loss of ACSS2 negatively impacts the transcription of activity-dependent genes required for LTM formation in a fear conditioning model. **a)** Volcano plot for gene expression changes between fear-conditioned WT and ACSS2KO mice 30 minutes after acquisition. (WT FC 30’: n=5; KO FC 30’: n=3, padj < 0.05). **b)** Volcano plot for gene expression changes between homecage-housed WT and ACSS2KO mice. (WT baseline: n=5; KO baseline: n=3, padj < 0.05). **c)** Top 10 GO terms (DAVID, BP) enriched in WT FC 30’ (DAVID, BP) compared to KO FC 30’. **d)** Top 10 GO terms enriched in KO FC 30’ (DAVID, BP) compared to WT FC 30’. **e)** Euler overlap of significantly altered genes in WT FC 30’ and KO FC 30’ relative to baseline controls (padj < 0.05) **f)** Bar plots of 12 well-characterized IEGs showing normalized transcript counts across all four conditions (RNA-seq; Deseq2). **g)** Heatmap of top 60 differentially expressed genes in WT FC 30’ mice compared to baseline showing sample-to-sample variability (RNA-seq; top 60 genes determined by padj value)

Instead of LTM-related genes that increase in the WT mouse following FC, the 1219 Acss2^KO^-biased genes showed a distinct focus on pathways downstream of transcription, with top gene ontology (GO) terms associated with protein translation (including ribosomal biogenesis and rRNA processing) (Fig. 4d). Many ribosomal proteins, including Rps5, Rps7, and Rps25, were highly upregulated in FC Acss2^KO^ mice compared to WT controls. The transcriptome of Acss2^KO^ mice also reflects increased metabolic needs of activated neurons, such as enrichment for key genes involved in every stage of the oxidative phosphorylation pathway, from Ndufa7 in complex I to Atp5e in complex V. However, consistent with previous results (Fig. 1f), no significant upregulation was observed in acetyl-coA-producing enzymes like Acly or Pdha1. In fact, transcription of Acly drops in the Acss2^KO^ hippocampus upon fear conditioning, agreeing with previous reports of its activity-dependent transcription^43^ (Supplementary Fig. 4f).

Given the number of genes dysregulated in ACSS2^KO^ after FC, we reasoned that some of the alterations could be constitutive changes in the mutant mouse and thus show differential expression under baseline conditions. Surprisingly, the hippocampal transcriptome of Acss2^KO^ mice was remarkably similar to WT mice without FC (Fig. 4b); by Deseq2 analysis, only three genes are differentially expressed at baseline, including Acss2, and Ifitm10 and Btaf1 barely surpassed the threshold for genome-wide significance (Fig 4b).

Since both Acss2^KO^ and WT mice were subjected to the same stimulus, and would thus be expected to upregulate a similar cohort of genes, we questioned how much of the dysregulation observed after FC (Fig 4a) was a blunting of activation in the KO animals, and how much was a discrete transcriptional response to stimulus between the genotypes. To determine how the dHPC transcriptome of WT and Acss2^KO^ mice responded to the stress of fear conditioning, we compared FC mice of each genotype to their respective baseline controls, then compared both sets of FC-specific transcripts to each other (Fig 4e). WT mice showed robust increase in expression of a subset of crucial activity-dependent genes called immediate early genes (IEGs), including Egr2, Egr1, Arc and Fos (Supplemental Fig. 4a). These genes are rapidly upregulated in response to neuronal activation, and include transcription factors and cytoskeletal proteins needed to initiate a cascade of activity-dependent transcription. This transcriptional program ultimately results in the stable synaptic remodeling that forms the molecular basis of LTM. The expression of IEGs in WT mice was reflected in a preponderance of GO terms related to DNA-templated gene transcription (Supplemental Fig. 4b). In contrast, while Acss2^KO^ mice upregulated a similar number of genes (n=705) as WT controls (n=652), IEGs were not strongly represented, while other transcriptional regulators, such as Arid4b, a subunit of the Sin3a corepressor complex, were among the most significantly upregulated genes (Supplemental Fig. 4c). As above, top GO terms show that Acss2^KO^ mice upregulated protein translation and mRNA processing, in clear contrast to the transcription-focused upregulation in WT mice (Supplemental Fig. 4d to Supplemental Fig. 4b). Interestingly, by adding the baseline conditions for each group, we saw that oxidative phosphorylation was downregulated upon FC in the WT mice, and that the Acss2^KO^ bias was due not to upregulation, but to a failure to similarly respond.

To determine the degree of similarity in FC-response between genotypes, we next examined the overlap between the two sets of FC-specific genes. We found that only 91 genes (7.1%) were shared between the two conditions. These 91 genes included transcriptional regulators (Arid4B, Btaf1, Smad5, ZFP369, Kdm3a, Med23, Pkn2) --including Creb1, a key transcription factor in the expression of IEGs. However, only three IEGs (Egr2, Arc, and Tiparp) were significantly upregulated in both genotypes (Fig. 4e). Plotting the relative expression of 12 well-characterized IEGs, we found that the induction of most was attenuated in Acss2^KO^ mice following fear conditioning. In all cases, the fold-change between baseline and fear conditioned Acss2^KO^ mice is less than in WT mice (Fig. 4f). A heatmap of the top 40 most differentially expressed genes between WT mice before and after fear conditioning (by adjusted p-value), showed a similar trend, where the expression of highly upregulated IEGs is dampened in Acss2^KO^ mice (Fig. 4g). These results indicate that ACSS2 is essential for the expression of most IEGs, and strongly suggest that LTM deficits in Acss2^KO^ mice are tied to reduced activity-dependent transcription. The striking contrast between ACSS2’s minimal impact on baseline expression and its significant impact on activity-dependent transcription may underlie the equivalent performance of ACSS2^KO^ mice to WT controls on LTM-independent behaviors (Supplementary Fig. 2), but the poor performance of ACSS2^KO^ on assays requiring long-term memory (Fig 1i, Fig. 2).

### Pharmacological disruption of ACSS2 in vitro

Our characterization of ACSS2^KO^ mice revealed a requirement for ACSS2 in fear memory consolidation. This finding led us to ask whether pharmacological inhibition of ACSS2 would similarly disrupt fear memory consolidation, as this could represent a potential therapeutic as a short-acting memory disruptor. We obtained a commercially available small molecule inhibitor of ACSS2 (referred to as cACSS2i) shown to be potent and specific in HepG2 cells, inhibiting the uptake of radiolabeled acetate into lipids and histones while not inhibiting enzymatic activity of related enzymes ACSF2 and ACSL5^23^. The effect of this ACSS2 inhibitor in the brain, however, is unknown.

We previously described that cACSS2i treatment in primary mouse hippocampal neurons growing in culture resulted in molecular phenotypes similar to genetic knockdown in a mouse neuronal cell line (mouse Cath. -a-differentiated CAD cells). These effects included reduced nuclear acetyl-CoA levels, lowered histone acetylation, lowered neuronal gene expression, and reduced biomarkers of neuronal cell differentiation^20^. Here, utilizing human pre-neuronal NT2 cells, which can be differentiated into neurons, we assessed histone acetylation following treatment with cACSS2i. Following treatment for 24hr, pre-neuronal NT2 cells were lysed and H3K9ac levels were examined by western blot, relative to total histone H3. Treatment with cACSS2i substantially reduced H3K9ac over a range of concentrations (Fig. 5a) (one-way ANOVA F (6,7) = 30.94, p=0.0001, *post hoc* Dunnett test comparing each concentration to DMSO adjusted p-value ≤ 0.0002). Thus, small molecule-mediated inhibition of ACSS2 in a human NT2 cell line reduces histone acetylation.

**Figure 5:**
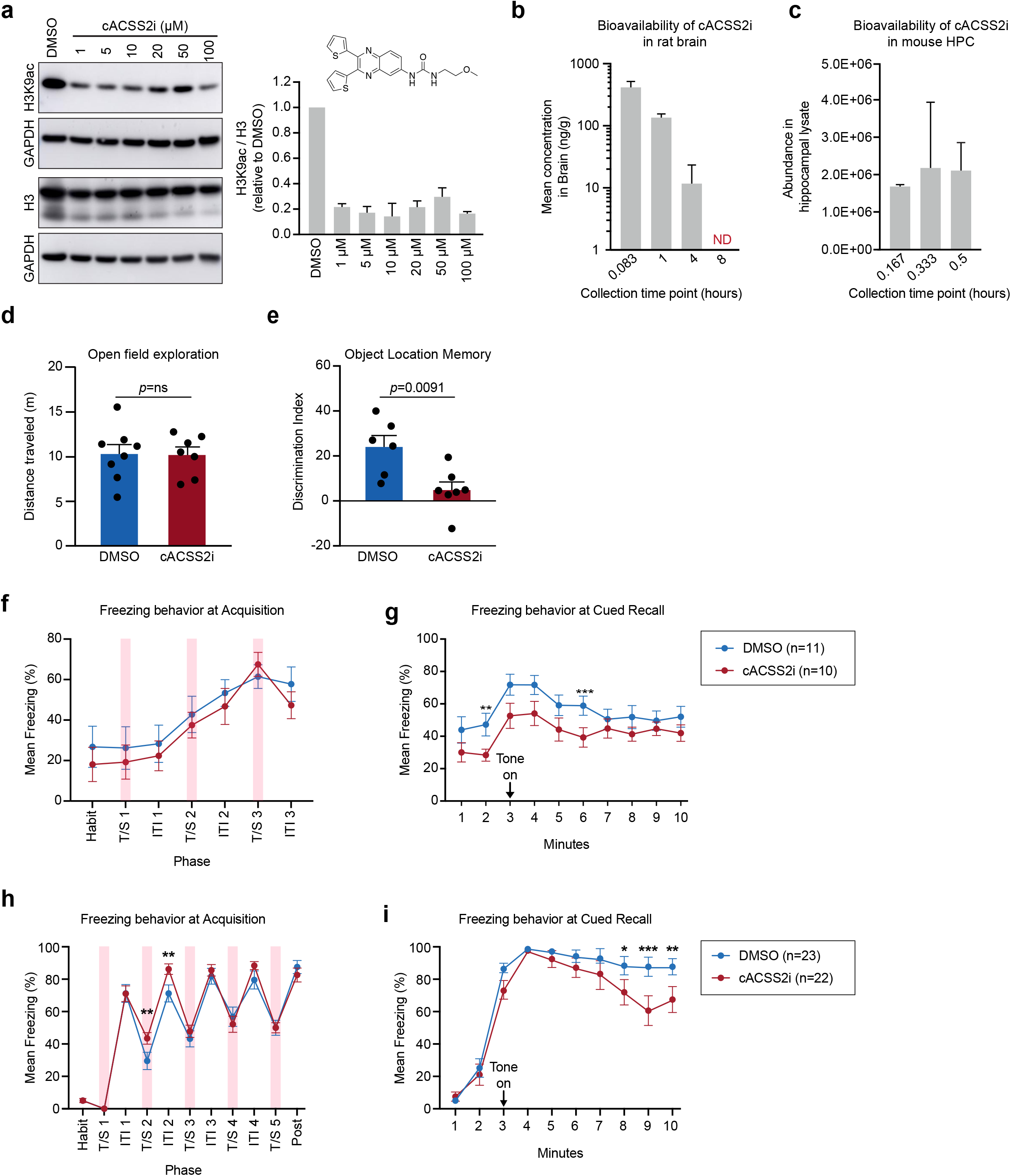
Small molecule inhibitor of ACSS2 reduces histone acetylation *in vitro*, is bioavailable in rodents, and disrupts long-term memory formation *in vivo*. a) Western blot on NT2 cell lysates for H3K9ac and H3 with respective GAPDH loading control following treatment with cASS2i. Quantification at right for two replicates shows a reduction in H3K9ac relative to total H3 (one-way ANOVA F (6,7) = 30.94, p=0.0001, *post hoc* Dunnett test comparing each concentration to DMSO adjusted p-value ≤ 0.0002). b) Pharmacokinetic analysis reveals presence of cACSS2i following IV administration in rat brain homogenate out to 4hr (n=3 animals at t0.083 and t1, n=1 at t4). cACSS2i is not detected at 8hr (ND). c) cACSS2i is detected in mouse hippocampal lysate at time points through 30 min following IP injection (n=2 for each time point). d) Open field assessment in mouse indicates no difference in basal locomotor activity following IP injection of DMSO vehicle or cACSS2i (DMSO n=8, cACSS2i n=7). e) Long term memory formation is disrupted in cACSS2i-injected animals in the object location memory assay (DMSO n=6, cACSS2i n=7, p=0.0091, unpaired student’s t-test). f) Freezing behavior in mouse during acquisition with cACSS2i treatment reflected as % time freezing/total time. Freezing during habituation period was averaged, remaining time is binned into 30sintervals. Timing of CS-US pairings indicated by transparent red bars (p =ns, two-way ANOVA). g) Freezing behavior during cued recall in mice treated with cACSS2i or DMSO at acquisition plotted as % time freezing/total time, binned into 30s intervals. Onset of tone is marked by a black arrow. Significant time points are observed at the onset of cue (two-way ANOVA, Fisher’s LSD **p=0.0308, ***p=0.0319). h) Freezing behavior in rat during acquisition with cACSS2i plotted as % time freezing/total time. Freezing during habituation period was averaged, remaining time is binned into 30s intervals. Time of CS-US pairings indicated by transparent red bars. Here, rats receiving cACSS2i exhibit small but significant increased freezing at the second CS-US presentation and during the second inter-trial-interval (two-way ANOVA, Fisher’s LSD **p=0.0375, 0.0208). i) Freezing behavior during cued recall in rats treated with cACSS2i or DMSO at acquisition plotted as % time freezing/total time, binned into 30s intervals. Onset of tone is marked by a black arrow. Significant time points are observed late in cue presentation (two-way ANOVA, Fisher’s LSD *p=0.0285, ***p=0.0003, *p=0.0070).

### Pharmacological inhibition of ACSS2 in vivo

Given that small molecule inhibitors of ACSS2 reduce histone acetylation *in vitro*, we investigated whether ACSS2i reduces long-term memory in mouse behavioral models similarly to genetic disruption of ACSS2 shown above. First, we examined the ability of ACSS2i small molecules to cross the blood-brain barrier. Pharmacokinetic assessment in rats revealed that cACSS2i was present in brain homogenate following intravenous (IV) administration of cACSS2i at 5 minutes, at 1 hour, and at 4 hours post administration, and levels dropped more than 100-fold at 4 hours (Fig. 5b). Importantly for potential therapeutic usage in which a temporally limited effect would be desired, cACSS2i was no longer detected in the brain at 8 hours after IV administration, with minimal cACSS2i detected in plasma 24 hours following IV administration) (Supplemental Fig. 5a). In addition, we observed similar kinetics for cACSS2i in mouse hippocampus following administration by intraperitoneal (IP) injection (Fig. 5c).

To examine the effect of cACSS2i on basal locomotor activity, we performed a 30-minute open-field assay. Mice injected with cACSS2i displayed equivalent locomotor activity to those injected with DMSO vehicle (p=0.949), indicating no deleterious effect on movement (Fig. 5d). As we previously reported for ACSS2 knock down^20^ and for Acss2^KO^ above (Figs. 1h, 1i), ACSS2 is required for hippocampal-dependent spatial learning in the Object Location Memory (OLM) paradigm. We thus tested whether cACSS2i is able to reduce long-term memory in this assay. In this OLM assay, we administered cACSS2i or vehicle by IP injection prior to and immediately following training sessions. Long-term memory was assessed 24h later, with one object moved to a different location. During recall, vehicle-injected mice exhibited increased exploration of the object moved to a novel location (Fig. 5e), while, in contrast, mice injected with cACSS2i displayed a reduced discrimination index relative to the control counterparts (Fig. 5e ; 24.01±5.07 vs 4.83±3.59, p=0.0091). Thus, cACSS2i administration reduces long-term memory in OLM as we previously observed utilizing genetic mutations of ACSS2, serving as proof-of-concept that pharmacological inhibition of ACSS2 is capable of disrupting memory consolidation.

### ACSS2i treatment reduces fear memory in a rodent cued fear conditioning model

We next investigated the ability of ACSS2i to disrupt fear memory consolidation in a rodent cued fear conditioning model as above and schematized in Supplemental Figure 6a. We administered cACSS2i via IP injection prior to and immediately following fear training. At 24 hours post-conditioning, mice were placed in an altered context and subjected to a singular extended tone in the absence of the foot shock. While DMSO-injected and cACSS2i-injected mice exhibited similar behavior during fear conditioning acquisition (Fig 5g), there was reduced freezing in cACSS2i mice during cued recall (Fig. 5h). These results were similar to the observations of the ACSS2^KO^.

We next tested fear memory in rats to increase our rigor through cross-species comparison. We examined whether cACSS2i could disrupt fear memory consolidation in a rat cued fear conditioning assay, similar to the mouse experiment above (Supplemental Figure 6b). After a habituation period, rats were exposed to the pairing of an auditory tone co-terminating with a mild foot shock. The pairing was repeated for a total of five presentations. We administered cACSS2i via IP injection prior to and immediately following fear training. At 24 hours post-conditioning, rats were placed in an altered context and subjected to a singular extended tone in the absence of the foot shock. While cACSS2i-injected rats exhibited slightly higher freezing at fear acquisition (Fig 5i), we observed reduced freezing in cACSS2i-injected rats during cued recall (Fig. 5j). Taken together, these data in mouse and rat indicate that animals receiving cACSS2i at training displayed reduced freezing during subsequent testing in response to the tone, suggesting that cACSS2i can disrupt fear memory consolidation *in vivo*.

### ACSS2i injection results in reduced generalized anxiety and cued recall in a rodent model of PTSD

To further elucidate the potential of ACSS2 inhibition as a therapeutic for PTSD, we utilized the Predator Scent Stress (PSS) model, which more thoroughly analyzes levels of specific and generalized anxiety following an aversive event.^44,45^. Briefly, rats were exposed to a predator scent or to a sham control with cACSS2i or DMSO vehicle administered through IP injections prior to and following exposure (Fig. 6a). Seven days later, generalized anxiety was assessed following PSS exposure through use of acoustic startle response (ASR) and elevated plus maze (EPM) assays. In controls, we observed a strong increase in startle amplitude in PSS-exposed animals relative to sham-PSS (Fig. 6b) (sham-PSS-DMSO 158.33±37.04 a.u., n=18 vs PSS-DMSO 500.55±48.55 a.u., n=20; Tukey’s multiple comparisons test p<0.0001). Startle amplitude was lower in cACSS2i-injeced animals exposed to PSS (Fig. 6b) (PSS-DMSO 500.55±48.55 a.u, n=20 vs 356.38±35.12 a.u, n=21; Tukey’s multiple comparisons test p<0.0001). PSS exposure also resulted in a significant increase in Anxiety Index in EPM activity, a metric integrating time spent in open arms, number of entries to open arms, and total exploration of the maze (Fig. 6c, Supplemental Figure 7) (sham-PSS-DMSO 0.72±0.03, n=18 vs PSS-DMSO 0.92±0.02, n=20; Tukey’s multiple comparisons test p<0.0001). Strikingly, this increase was blunted in cACSS2i-injected PSS rats relative to DMSO-injected animals (Fig. 6c) (PSS-DMSO 0.92±0.02, n=20 vs. PSS-cACSS2i 0.75±0.03, n=21; Tukey’s multiple comparisons test p<0.0001). Importantly, all cohorts displayed similar levels of activity in EPM, indicating that the observed differences were due to variance in anxiety level rather than a disruption of locomotor activity (Fig 6d). Taken together, EPM and ASR data indicate that cACSS2i administration reduces fear and generalized anxiety behavior in rat models following exposure to PSS.

**Figure 6:**
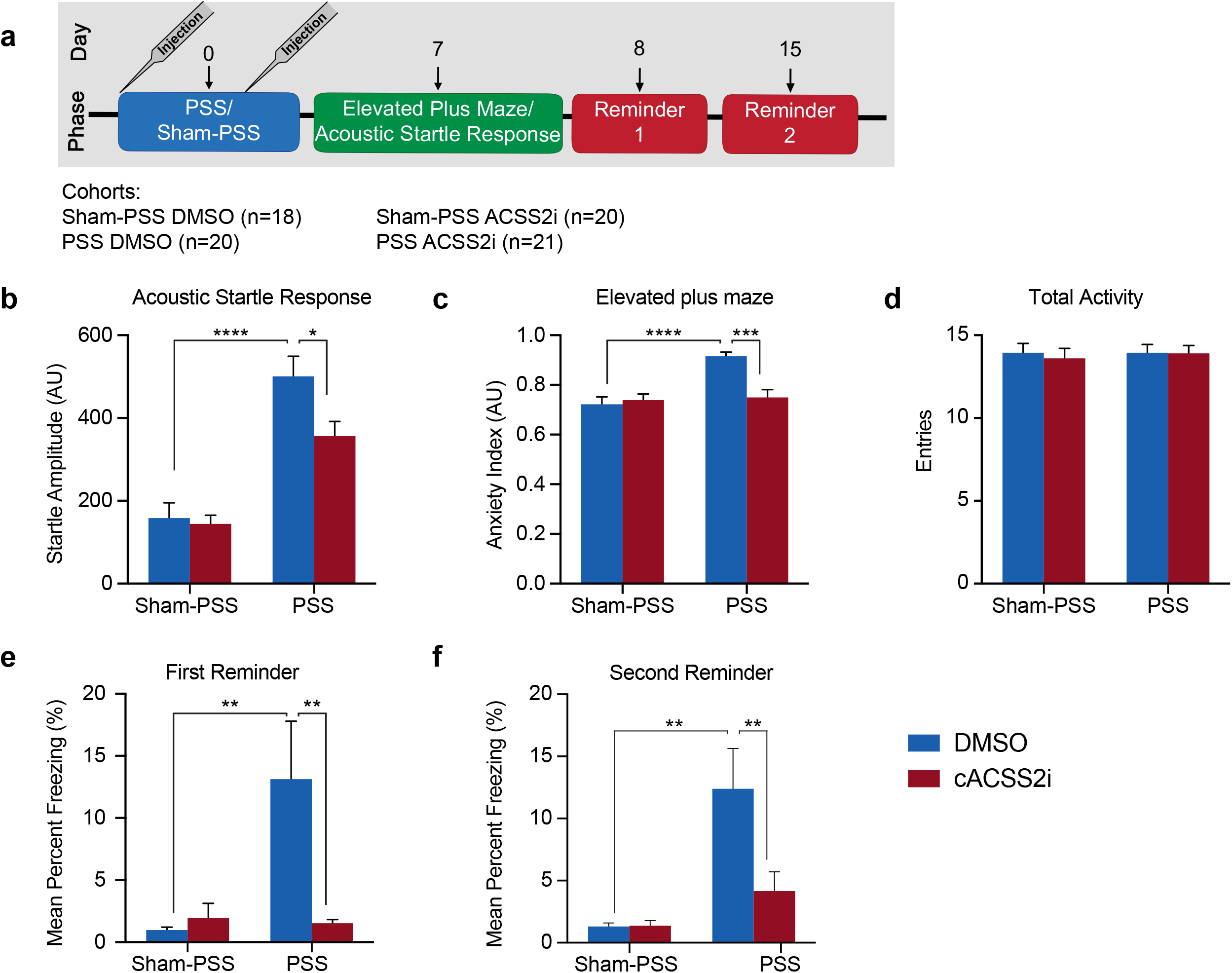
cACSS2i treatment impairs predator scent stress-induced fear memory consolidation in rats. a) Outline of the PSS assay. Rats are exposed to PSS (soiled cat litter) or sham-PSS (fresh cat litter) with cACSS2i or DMSO on board. One week later, anxiety level in animals is measured through elevated plus maze and acoustic startle response. The following day, animals are re-exposed to the cat litter stimulus and freezing behavior is measured. A second reminder exposure takes place one week later. b) PSS-exposed DMSO-injected rats display increased startle amplitude relative to sham-exposed DMSO-injected rats. cACSS2i-treated PSS-exposed rats exhibit a reduced startle amplitude. c) PSS-exposed DMSO-injected rats display increased Anxiety Index relative to sham-exposed DMSO-injected rats. cACSS2i-treated PSS-exposed rats exhibit a lower Anxiety Index. d) Total entries into open and closed arms of the EPM apparatus to indicate total activity in the assay. No differences between treatment groups were detected. e) PSS-exposed DMSO-injected rats display increased freezing relative to sham-exposed DMSO-injected rats at the first reminder session. No such increase is observed in cACSS2i-treated PSS-exposed rats. f) The effects observed at the first reminder session persist 1 week later at a second reminder session. PSS-exposed DMSO-injected rats display increased freezing relative to sham-exposed DMSO-injected rats at the first reminder session. No such increase is observed in cACSS2i-treated PSS-exposed rats.

One day following assay of anxiety behavior (Day 8), rats were re-exposed to the cat litter stimulus in a similar environment and freezing behavior was measured. PSS-exposed DMSO-injected rats exhibited significant increase in freezing during the reminder session compared to Sham-PSS DMSO-injected animals (sham-PSS-DMSO 0.98±0.23, n=18 vs PSS-DMSO 13.13±4.65, n=20; Tukey’s multiple comparisons test p=0.005). In contrast, cACSS2i-injected PSS rats exhibited significantly lower freezing than DMSO-injected PSS animals (Fig. 6e), on par with sham-PSS treated rats, indicating reduced fear memory. Finally, we assessed long-term retention of the stress memory in a second recall session seven days following the first reminder session (Day 15). PSS-exposed rats injected with DMSO continued to display significant freezing relative to Sham-PSS controls. Again, cACSS2i-injected PSS-exposed rats displayed decreased freezing (Fig. 6f), indicating that reduction in fear memory persisted in cACSS2i-injected PSS-exposed rats. These results thus suggest that inhibition of ACSS2 disrupts long-term memory consolidation surrounding trauma exposure.

### Development of a novel inhibitor of ACSS2

These findings with cACSS2i led us to design and synthesize novel analogs of ACSS2i in an effort to further improve the window of brain bioavailability and to optimize pharmacokinetics for acute and limited disruption of memory formation. Toward that end, we replaced the methoxyethyl moiety present in ACSS2i with an n-pentyl group to generate nACSS2i, with increased lipophilicity and therefore increased blood-brain barrier penetration^46^ The new analog was prepared from the isocyanate (triphosgene, Hunig’ base, dichloromethane) derived from 2,3-di(thiophen-2-yl)quinoxalin-6-amine^47^. This novel compound (hereafter nACSS2i) was tested in histone acetylation *in vitro*, and as with the cACSS2i, we observed significant reduction in levels of H3K9 acetylation following treatment with nACSS2i (Fig. 7a; One-way ANOVA F (6,7) = 17.38, p=0.0007, *post hoc* Dunnett each concentration compared to DMSO adjusted p-value ≤ 0.0013). Further, we detected nACSS2i in rat plasma and in brain homogenate following IV administration (Fig. 7b, Supplemental Fig. 5b). Intriguingly, nACSS2i may be cleared faster from the brain than cACSS2i, as nACSS2i was not detected at 4 hours post-administration.

**Figure 7:**
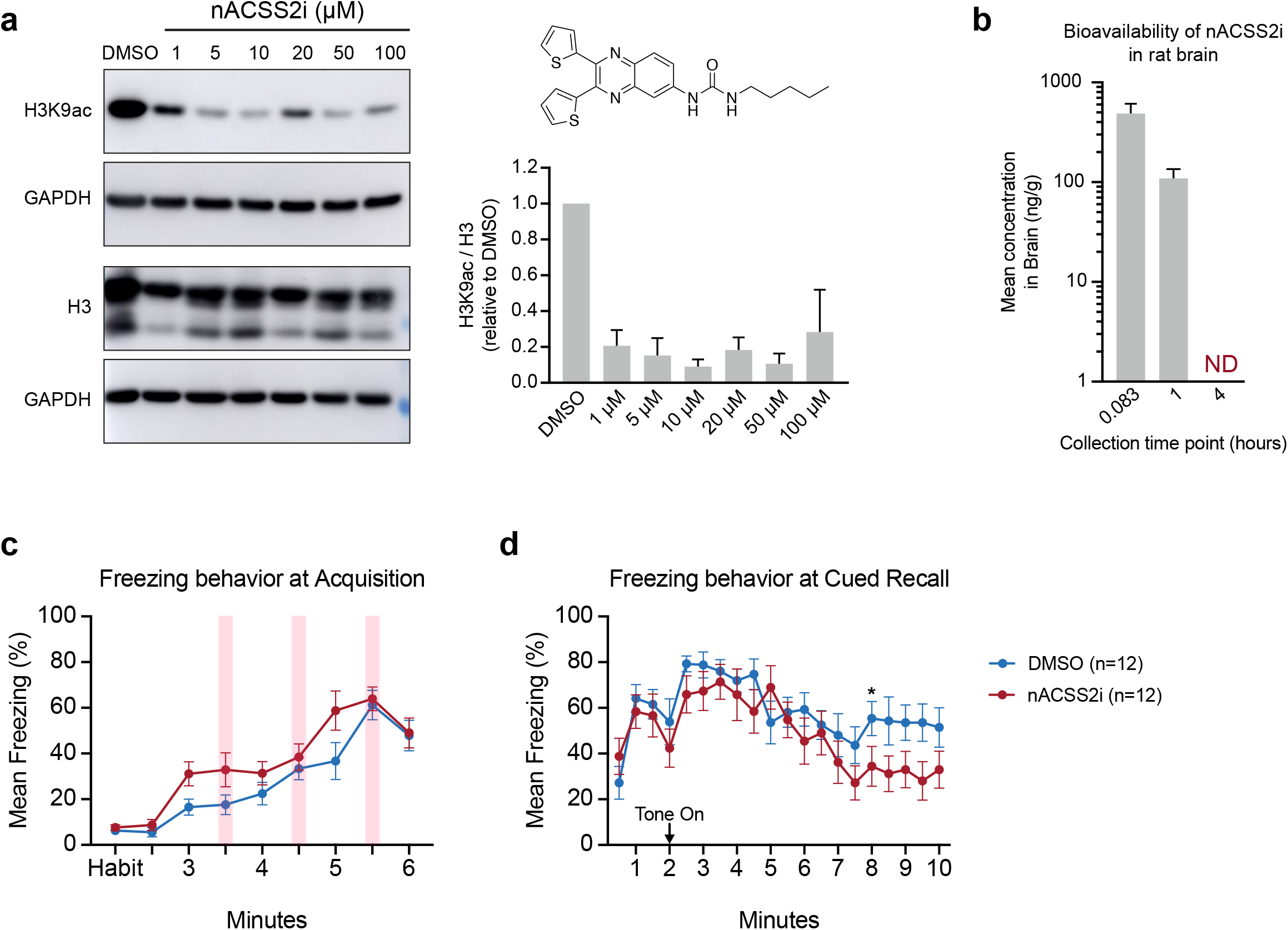
A novel small molecule inhibitor of ACSS2 reduces histone acetylation in vitro, is bioavailable in rodents, and disrupts long-term memory formation in vivo. a) Western blot on NT2 cell lysates for H3K9ac and H3 with respective GAPDH loading control following treatment with nASS2i. Quantification at right for two replicates shows a reduction in H3K9ac relative to total H3 (One-way ANOVA F (6,7) = 17.38, p=0.0007, post hoc Dunnett each concentration compared to DMSO adjusted p-value ≤ 0.0013). b) Pharmacokinetic analysis reveals presence of nACSS2i following IV administration in rat brain homogenate out to 1hr (n=3). c) Freezing behavior in mouse during acquisition with nACSS2i treatment reflected as % time freezing/total time. Freezing during habituation period was averaged, remaining time is binned into 30sintervals. Time of CS-US pairings indicated by transparent red bars (p =ns, two-way ANOVA). d) Freezing behavior during cued recall in mice treated with nACSS2i or DMSO at acquisition plotted as % time freezing/to-tal time, binned into 30s intervals. Onset of tone is marked by a black arrow. Significant time points are observed late in cue presentation (two-way ANOVA, Fisher’s LSD **p=0.0408).

To examine whether nACSS2i was similarly able to disrupt fear memory consolidation, mice were subjected to cued fear conditioning as described above. While DMSO-injected and nACSS2i-injected mice exhibited equivalent behavior during fear conditioning acquisition (Fig. 7c), we observed reduced freezing in nACSS2i mice during cued recall. This reduction in freezing behavior occurred later in the onset of the tone relative to the phenotype observed with cACSS2i treatment (Fig. 7d; p=0.0408; Fisher’s LSD). Thus, pharmacological inhibition of ACSS2 with this novel small molecule may represent a promising therapeutic for PTSD.

## DISCUSSION

This study demonstrates that ACSS2 is a critical regulator of fear memory formation in mice and in rats. Both genetic ablation and small molecule inhibition of ACSS2 were sufficient to attenuate the consolidation of long-term fear memories. Importantly, loss of ACSS2 is well-tolerated. Mice harboring a whole body knockout of Acss2 were indistinguishable from WT mice in working memory tests and other LTM-independent tasks. Histone analysis of mice undergoing active consolidation of fear memories showed significant decreases in the histone marks H4K5ac and H3K9ac in Acss2^KO^ mice, with H4K8ac and H4K12ac showing a similar downwards trend compared to WT controls. This loss of histone acetylation was specific to the fear conditioned state, as untrained homecage-housed WT and untrained Acss2^KO^ mice showed similar levels of acetylation. RNA-seq analysis underscored the requirement for Acss2 during fear memory consolidation: at baseline, we observed only three differentially expressed genes in the dHPC; 30 minutes after stimulation, this number increased dramatically to 2873. Consistent with these results, we found that administering a small molecule inhibitor of ACSS2 (ACSS2i) during fear memory consolidation attenuated the persistence of fear memory in both mice and rats. Together, these genetic and pharmacological approaches show that ACSS2 is specifically involved in long-term memory and point to ACSS2 as a promising therapeutic target for treating pathological fear memories which underlie trauma, such as in PTSD.

Our results are consistent with previous findings that histone acetylation is a critical regulator of fear memory formation in the dHPC. Many histone residues are acetylated upon fear conditioning, including H3K9ac, H3K14ac, H4K5ac, H4K8ac, and H4K12ac^43^. Reductions of the histone acetyltransferases CBP, p300, and Kat2a in the dorsal hippocampus or the amygdala negatively impact the formation of cued and contextual fear conditioning^34,40–42,48^, as does the inhibition of another histone acetyltransferase (PCAF/Kat2b) in the prelimbic cortex^49^. Conversely, both cued and contextual fear conditioning are strengthened by the ablation of the histone deacetylases HDAC2 and Sirt1^50,51^. Systemic administration of chemical inhibitors of CBP (garcinol^52^) and HDACs (SAHA, Sodium Butyrate, Valproic acid, TSA)^6,50,53–55^ have been well studied, with CBP inhibitors administered during consolidation reducing fear memories and HDAC inhibitors enhancing them. Our results show that both Acss2 genetic ablation by germline knock out and ACSS2 inhibition by small molecule inhibitor reduced fear memory and histone acetylation similarly to CBP inhibition. However, importantly, while genetic knock out of CBP or p300 causes embryonic inviability^56,57^, the Acss2^KO^ is well-tolerated.

We found that reductions of H3K9ac and H4K5ac during fear conditioning coincided with the downregulation of many genes essential for LTM formation. In particular, we found that expression of key immediate early genes (IEGs) (including Egr1, Nr4a1, and Fosb) was attenuated in fear-conditioned Acss2^KO^ mice compared to WT controls. This is consistent with CBP’s contribution to the expression of many IEGs, including Fos^34^, Fosb^58^, Junb, and Nr4a1^59^. It is also consistent with observations that HDACs, especially HDAC2 and HDAC3, play critical roles in the regulation of Egr1, Fos^50^, Arc, and Nr4a1^60^.

The observed decrease of histone acetylation in the dHPC of fear-conditioned ACSS2^KO^ mice is likely due to reduced acetyl-coA availability. However, RNA-seq analysis revealed that loss of Acss2 also reduces the expression of key histone acetyltransferases, including CBP/Crebbp, p300, and Kat2a/Gcn5 (Supplemental Fig. 4e), suggesting a feed-forward loop caused by Acss2^KO^. The histone targets of these HATs include both of the most severely affected residues, H3K9ac and H4K5ac. H3K9ac is deposited mainly by Gcn5^61^, and H4K5ac by multiple HATs, including CBP/p300^62^. In turn, CBP and p300 are regulated by the immediate early gene Egr1^63^, which is severely attenuated in the Acss2^KO^ mouse after fear conditioning, suggesting that the dysregulation of epigenetic enzymes could in part be a downstream consequence of Acss2 manipulation, further affecting histone acetylation. Of note, the interconnected gene regulatory networks underlying LTM formation necessitated examining Acss2 at the earliest stages of LTM consolidation – beyond the expression of IEGs, it becomes difficult to determine which activity-dependent changes can be tied directly to ACSS2, and which are simply a function of reduced IEG expression.

Also arguing for specific effects of ACSS2 on long-term memory, we found that constitutive loss of Acss2 did not impact global histone acetylation levels in the dHPC. By histone mass spectrometry, we observed no difference between homecaged-housed Acss2^KO^ mice and WT controls (Fig 3). In *S. cerevisiae*, by contrast, loss of the Acss2 homolog Acs2 results in global histone deacetylation and transcriptional repression^17^. This reflects the importance of acetate in yeast metabolism; in the mammalian brain, glucose is the most important bioenergetic substrate. Therefore, in Acss2^KO^ mice, the existing histone acetylation present at baseline is likely provided by ACLY, which converts glucose-derived citrate into acetyl-coA, and is known to play an essential role in histone acetylation^18^. Though levels of Acly did not increase in compensation for Acss2 loss, the baseline RNA expression level of Acly is twice that of Acss2 in mouse hippocampus (compare Figs. 1g, 1f). Importantly, the striking differences seen in histone acetylation and transcription after neuronal stimulation point to a unique role for Acss2 during periods of extreme metabolic requirement, like neuronal stimulation^64^ — a process for which Acly does not appear to compensate.

Maintaining a balance of histone acetylation levels has been extensively studied and is critical for memory and cognition processes. Moreover, the disruption of these levels is disrupted in pathogenic states such as Alzheimer’s disease (AD)^65–67^. In an AD transgenic mouse model, phenylbutyrate improves levels of histone acetylation in while reducing Tau pathology^68^. Treatment with HDAC inhibitor vorinostat improves spatial memory in another AD mouse model^69^. While much promise surrounds the use of HDAC inhibitors in the clinic, persistent use of HDAC inhibitors also carries potential risks with toxicity and lack of specificity. Thus, development of inhibitors to ACSS2 with temporally limited effects may provide a promising alternative path to treating neuropsychiatric disorders.

In order to ensure the viability of ACSS2 to treat memory-related disorders, we focused on improving the ability of ACSS2i to cross the blood-brain barrier (BBB), an exquisitely selective network of epithelial cells forming tight junctions. The BBB serves as a significant hurdle in developing therapeutics for neurological disorders, acting as a barrier to entry to an estimated 98% of small molecule drugs^70,71^. Crucially, our *in vivo* pharmacokinetic data demonstrated that the ACSS2i is able to cross the blood-brain barrier. Through structure activity relationship studies to optimize BBB crossing, we developed a novel ACSS2i (nACSS2i) with altered lipophilicity^46^. Furthermore, a distinguishing feature in our drug development process was the temporal considerations of inhibiting ACSS2. That is, unlike more standard drug development approaches, our medicinal chemistry was driven by optimization of a temporally limited compound with a short half-life, as long-term memory consolidation requires rapid increases in histone acetylation and gene transcription in a sensitive time window, and additionally, with desirable rapid metabolism of the inhibitor. Thus, the transient nature of nACSS2i will reduce potential off-target effects associated with use of a memory disrupting drug.

We also demonstrated that these inhibitors reduce anxiety in a predator scent stress (PSS) model, which recapitulates heightened anxiety level and trigger-induced episodes present in PTSD^72^. After an aversive presentation of predator scent (PS), rats injected with ACSS2i during the consolidation window not only exhibited reduced freezing in response to PS-specific reminders, but also showed reduced anxiety in acoustic startle and elevated-plus maze assays. This striking ability of ACSS2i to disrupt fear memory consolidation and associated anxiety (Fig 6) again highlights the need to limit the temporal effects of this small molecule inhibitor, so as not to interfere with general memory formation. Thus, taken together, the specificity of ACSS2 towards rapid activity-dependent histone acetylation and gene transcription makes it an attractive therapeutic target in the treatment of diseases of unwanted memory, including PTSD.

## MATERIALS AND METHODS

### Mouse experiments

Animal use and all experiments performed were approved by the Institutional Animal Care and Use Committee (IACUC, protocols 804849 and 804484). All personnel involved have been adequately trained and are qualified according to the Animal Welfare Act (AWA) and the Public Health Service (PHS) policy. Experiments utilizing ACSS2 inhibitors were performed in 8-12 wk old male mice. For experiments involving Acss2^KO^ mice, 4-month old adult male mice were used. To prevent genetic drift, Acss2^KO^ and corresponding WT controls were generated through homozygous crosses of F1 progeny from heterozygote breeding cages. C57Bl6/J mice for the inhibitor assays were purchased from Jackson Laboratories. Mice were housed on a 12/12h light/dark cycle (7 am to 7 pm), with food and water provided ad libitum. All behavioral experiments were conducted between 7 am and 11 am to reduce time-of-day effects.

### Generation of Acss2 knockout mice using CRISPR-Cas9

We used CRISPR/Cas9 technology to generate constitutive Acss2^KO^ mice as previously described^73^. Pairs of single guide RNAs (sgRNA) targeting intronic regions of the *Acss2* locus upstream of exon 3 and downstream of exon 7 were designed with the CRISPOR online tool^74^ (http://crispor.org) using the mm10 genome assembly as template. The resulting guides (5’[exon 3]: were selected to maximize specificity, while retaining high on-target efficiency. sgRNAs were synthesized by *in vitro* transcription (T7 High Yield RNA Synthesis Kit, NEB, E2040S). Cas9 mRNA (100 ng/uL. Trilink, L-6125) and both sgRNAs (100 ng/uL each) were diluted in injection buffer (10 mM Tris, pH 7.5; 0.1 mM EDTA). This mix was injected into the cytoplasm of single-cell C57Bl6/J mouse embryos. Resulting founder mice were screened for the desired deletion using 5 PRIME Hot Master Mix (Quantabio, 2200400) and primer pairs designed to amplify across each cut site: (5’: F-TGGCCTCCAACACTCTAAGT; R - CTGTGCCCGCTCAACATATG ; 3’: R - GAATCCCTTTTCTGCGTCCC), and flanking the entire deletion (F-TGGCCTCCAACACTCTAAGT; R – GAATCCCTTTTCTGCGTCCC). Sanger sequencing was used to confirm sites of non-homologous end joining. To reduce the risk of off-target effects, founder mice were backcrossed to C57Bl6/J mice (obtained from Jackson Laboratories) for three generations. Experimental mice were genotyped in a single reaction (ACSS2 F-TGGCCTCCAACACTCTAAGT; ACSS2 R1-CTGTGCCCGCTCAACATATG; ACSS2 R2-GAATCCCTTTTCTGCGTCCC).

### General Synthesis of ACSS2 Inhibitors

2,3-di(thiophen-2-yl)quinoxalin-6-amine was dissolved in anhydrous dichloromethane and Hünig’s base was added. A solution of triphosgene in dichloromethane was then added and the reaction was stirred for four hours at 25°C. The corresponding amine was added and the reaction was stirred for sixteen hours at 25°C. The reaction was then concentrated and the resulting residue was purified by silica gel chromatography to afford the inhibitor analogs.

### Acss2i administration

Acss2 inhibitors were dissolved in an injection vehicle containing 30% (w/v) 2-Hydroxylpropyl-beta-cyclodextrin (HP-β-CD), and 1-5% DMSO. For certain experiments (Fig 5d, f, g) 100 mM acetate was included in the vehicle as a buffering agent. Dosages of 4 mg/kg were delivered via intraperitoneal injection.

### Open field assay

Mice were placed in an open arena (30 cm x 40 cm) and allowed to explore freely for 6 minutes. Resulting videos were analyzed using ANY-maze™ behavioural tracking software (Stoelting Co.). Locomotion was assayed using path length (in meters) over the entire 6 minute period. Thigmotaxis (a measure of anxiety) was measured by dividing the amount of time spent within the peripheral zone (defined as 8 cm from the arena walls) over the entire period.

### Y-maze test of spontaneous alternation

The Y-maze test of spontaneous alternation assess working spatial memory in rodents^28^. Mice were gently placed in the distal portion of an arm in a Y-shaped arena and allowed to explore freely. Mice were recorded by video cameras, and arm entries were scored by eye. Arm entries were called when the mouse’s hindquarters passed fully into the new arm. Percent spontaneous alternations were quantified according to the formula [SAR = [Number of alternations/(total arm entries – 2)] x 100.

### Acoustic startle response

Mice were placed in acoustic startle chambers (SR-Labs). Broadband acoustic startle bursts were emitted from a high-frequency speaker mounted above the mouse chamber. To generate input/output response curves, randomized bursts of various dB levels (100 db, 110 db, 120 db, 6 times each) were presented. The response to discrete stimuli was reported as a change in current (Vmax) measured by a voltmeter. Startle responses were measured by a Piezo Electronics monitor mounted under the stage platforms.

### Object location memory paradigm

Non-aversive spatial memory was assessed as previously described^20^, with minor modifications to accommodate dosing with the ACSS2 inhibitors (ACSS2i). Briefly, mice were handled for 2 minutes a day for three consecutive days (KO), or (x days – inhibitor). During handling, all mice used in ACSS2i experiments were injected intraperitoneally with PBS. On training day, mice were habituated to an empty arena for 6 minutes, then exposed to a collection of three distinct objects (glass bottle, metal tower, plastic cylinder) of roughly similar sizes for 6 minutes. This training was repeated three times, with mice removed and all surfaces wiped down with 70% ethanol between trainings. For testing, the mice were returned to the same arena 24 hours later, this time with one object moved to a new location. Mice were allowed to explore freely for 6 minutes. Mice were monitored using a video camera, and time spent interacting with each object was assessed afterwards. Quantification of mouse-object interactions was blinded. Discrimination index was calculated according to [DI = (t_displaced_ – [(t_stationary1_ + t_stationary2_)/2])/(t_displaced_ + [(t_stationary1_ + t_stationary2_)/2) × 100]]

### Cued and contextual fear conditioning (mice)

Mice were handled for three consecutive days for 2 minutes each prior to the start of the experiment. Mock injections were performed on subjects of ACSS2i assays. On the day of fear acquisition, mice were individually placed in conditioning chambers (Med Associates) and habituated to the novel environment for 2 minutes. An auditory cue (an un-modulating tone: 80 dB and 5 kHz) was presented for 30 seconds, co-terminating with a mild 2s, 1 mA foot shock. ACSS2i was administered by IP injection (ACSS2i in 25%HBCD in PBS, 1%DMSO, 4mg/kg) 5 minutes prior to being placed in the fear conditioning chamber and 30 minutes after being removed from the chamber. Chambers were wiped down with 70% ethanol between each round. After an inter-tone interval of 1 minute, the tone-shock pairing was presented twice more (for a schematic, see Fig. 2a). Mice were promptly removed 30s after shock onset and returned to their home cages. To assess retention of LTM related to the auditory cue, chambers were modified to remove spatial and olfactory cues. The shape of the chamber was modified with cardboard inserts, and the barred flooring replaced with a solid layer. The scent of ethanol was masked with vanilla extract. The same tone (minus the shock) was repeated either in the same pattern as the previous day (Acss2^KO^ experiments) or for a continuous 7 minutes (Acss2i experiments). Freezing behavior was monitored automatically using FreezeScan™ software (CleverSys, Inc) for the entire recall period. For the single shock contextual fear conditioning paradigm, mice were habituated to the chamber for 2.5 minutes prior to the administration of a mild 2s, 1.5 mA foot shock. 24 hours later, freezing response was tested for 5 consecutive minutes.

### Cued fear conditioning (rats)

Male Sprague-Dawley rats, aged 7 to 9 weeks, were handled for three consecutive days for 2 minutes each prior to the start of the experiment. On the day of fear acquisition, rats were acclimated to the procedure room for 30 minutes. ACSS2i was administered by IP injection (ACSS2i in 5%DMSO in 0.5% methylcellulose at 4mg/kg) 5 min prior to being placed in the fear conditioning chamber and 30 minutes after being removed from the chamber. Automated chambers (Kinder Scientific, Poway CA) using infrared beams to detect movement were used for the assay. An almond scent was place under the grid floor during the entire session. Fear training consisted of a 3-minute habituation followed by a 20-second, 80dB tone; during the last 3 seconds of the tone animals received a 1.5 mA foot shock. This pairing was repeated twice at 1-minute intervals for a total of five paired presentations of tone and shock. Freezing behavior was be recorded in 10-sec intervals during the session. To assess cued fear memory 24 hours later, rats were placed in the chambers with a different (lemon) scent and black Plexiglas floor over the grid floor. Freezing behavior in response to an altered context was recorded for 2 minutes in 10-sec intervals. The 80dB tone was be presented for 8 minutes and immobility recorded in 10-sec intervals to measure freezing response to the tone cue.

### Predator Scent Stress (PSS)

Male Sprague-Dawley rats (between 190-220 g) were individually placed on well-soiled cat litter, which was used by a cat for 2 days and sifted for stools. They were exposed to the litter for 10 min in a plastic cage (inescapable exposure) placed on a yard paving stone in a closed environment. Sham-PSS was administered under similar conditions, but the rats were exposed to a fresh, unused cat litter.

### Behavioral measurements

The behavior of rats was assessed in the EPM and ASR paradigms, as described previously^72,75^ and as briefly detailed below.

### Elevated plus-maze

The maze was a plus-shaped platform with two opposing open arms and two opposing closed arms (closed arms surrounded by 14-cm high opaque walls on three sides)^76^. Rats were placed on the central platform, facing an open arm, and were allowed to freely explore the maze for 5 minutes. Each test was videotaped and the behavior of the rat was subsequently scored by an independent observer. Arm entry was defined as entering an arm with all four paws. At the end of the 5 min test period, the rat was removed from the maze, the floor was wiped with a damp cloth, and any faecal boluses removed.

The percentage of time spent in the open arms [time spent in open arms/(time spent in open arms + time spent in closed arms)×100] and the percentage of the number of open-arm entries [open arms entries/(open arms entries +closed arms entries)×100] was used as a measure of anxiety.

### Acoustic startle response

Startle responses were measured by using two ventilated startle chambers (SR-LAB system, San Diego Instruments, San Diego, CA). The SR-LAB calibration unit was used routinely to ensure consistent stabilimeter sensitivity between test chambers and over time. Each Plexiglas cylinder rested on a platform inside a soundproof, ventilated chamber. Movement inside the cylinder was detected by a piezoelectric accelerometer below the frame. Sound levels within each test chamber were measured routinely with a sound level meter (Radio Shack) to ensure consistent presentation. Each test session began with a 5-min acclimatization period to background white noise of 68 dB, followed by 30 acoustic startle trial stimuli presented in 6 blocks (110 dB white noise of 40 ms duration and 30 or 45 sec inter-trial interval). Two behavioral parameters were assessed: (a) the mean startle amplitude (averaged over all 30 trials); and (b) the percent of startle habituation to repeated presentation of the acoustic pulse. For the latter, the percent change was calculated between the response to the first and last (6^th^) blocks of sound stimuli, as follows:

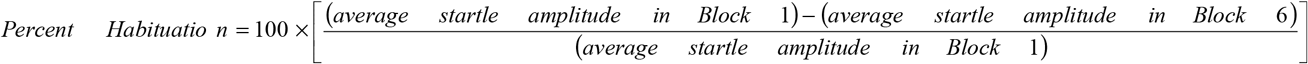

### RNA extraction and RNA-seq library preparation

RNA was extracted and libraries were prepared as previously described^21^. Briefly, total RNA was extracted using Trizol-chloroform from either whole HPC (Fig 1, Supplementary Fig. 1) or 1 mm-thick slices of dHPC (comprising the CA1, CA2, CA3, and dentate gyrus). Total RNA quality was assessed on the Bioanalyzer platform using the RNA 6000 Nano assay (Agilent). mRNA was isolated from 300 ng total RNA using the NEBNext® Poly(A) mRNA Magnetic Isolation Module (E7490L), and libraries were prepared using the NEBNext® Ultra™ II RNA Library Prep Kit for Illumina® (E7770). All RNA-seq data were prepared for analysis as follows: NextSeq sequencing data was demultiplexed using native applications on BaseSpace. Demultiplexed FASTQs were aligned by RNA-STAR 2.5.2 to assembly mm10 (GRCm38). Aligned reads were mapped to genomic features using HTSeq 0.9.1. Quantification, library size adjustment, and differential gene expression analysis was done using DESeq2. Gene list overlaps were tested for significance using the hypergeometric test.

### Western blots

Cells and tissue were lysed in RIPA buffer containing 50□mM Tris pH 8.0, 0.5□mM EDTA, 150□mM NaCl, 1% NP40, 1% SDS, supplemented with HALT protease and phosphatase inhibitor cocktail (Life Technologies, number 78446) and sodium butyrate (10 mM). Protein concentration was determined by BCA protein assay (Life Technologies, number 23227), and equal amounts of protein were loaded onto 4–12% Bis-Tris polyacrylamide gels (NuPAGE). Proteins were transferred to nitrocellulose membranes and subsequently blocked with 5% milk in TBS-T (blocking buffer). Membranes were incubated with primary antibodies diluted in blocking buffer for 4□°C overnight. Membranes were washed three times in TBS-T for 10 minutes each before incubation with HRP-conjugated secondary antibodies in blocking buffer. Membranes were washed again as before, developed with SuperSignal west pico PLUS chemiluminescent substrate (Thermo Fisher) then imaged with an Amersham Imager 600.

### Antibodies

The antibodies used were anti-H3 (Abcam ab1791), anti-H3K9ac (Abcam ab4441), anti-ACSS2 (CST 3658) anti-GAPDH (CST 5174).

### Histone extraction, propionylation, and digestion

Histones were extracted from 1 mm^2^ punches taken from the dorsal CA1 region of the mouse hippocampus by using Nuclei Isolation Buffer (NIB) as previously described^33^. The tissue punches were incubated in NIB (15 mM Tris-HCl, 15 mM NaCl, 60 mM KCl, 5 mM MgCl_2_, 1 mM CaCl_2_ and 250 mM sucrose at pH 7.5; 0.5 mM AEBSF, 10 mM sodium butyrate, 5 μM microcystin and 1 mM DTT added fresh) with 0.2% NP-40 on ice for 5 min. Two rounds of NIB incubation were performed at a volume buffer:cell pellet of 10:1; the first round 0.2% NP-40 was added to lyse the cell membrane, and the second without NP-40 to remove the detergent from the nuclear pellet. Each step included centrifugation at 700 x g for 5□min to pellet the intact nuclei. Next, the pellet was incubated in 0.2□M H_2_SO_4_ for 2□hours, and the supernatant was collected after centrifugation for 5□min at 3400 x g. Finally, histones were precipitated with 33% trichloroacetic acid (TCA) overnight. The histone pellet was then washed with ice-cold acetone to remove residual TCA.

Histones were derivatized and digested as previously described^33^. Histone pellets were resuspended in 20 μl of 50 mM ammonium bicarbonate (pH 8.0), and 10 μl derivatization mix was added to the samples, which consist of propionic anhydride and acetonitrile in a 1:3 ratio (v/v), and this was immediately followed by the addition of 5 μl ammonium hydroxide to maintain pH 8.0. The sample was incubated for 15 min at 37□°C, dried and the derivatization procedure was repeated one more time to ensure complete derivatization of unmodified and monomethylated lysine residues. Samples were then resuspended in 50 μl of 50 mM ammonium bicarbonate and incubated with trypsin (enzyme:sample ratio of 1:20) overnight at room temperature. After digestion, the derivatization reaction was performed again twice to derivatize the N-termini of the peptides. Samples were desalted using C_18_ stage tips before LC–MS analysis and dried. Finally, the peptides were resuspended in 0.1% formic acid (FA) prior to nLC-MS/MS.

### Nano LC-MS/MS for bottom-up histone peptide analysis

The peptide mixture was separated using a Dionex Ultimate 3000 high-performance liquid chromatography (HPLC) system with a two-column setup, consisting of a reversed-phase trap column (Acclaim PepMap100 C18; Thermo Fisher Scientific) (5-μm pore size, 100□Å, 300-μm inner diameter [i.d.] by 5□mm) and a reversed-phase analytical column (30□cm, 75-μm i.d., 360□μm o.d., packed in-house with Pur C18 AQ [Dr Maisch; 3-μm pore size]). The loading buffer used was 0.1% trifluoroacetic acid (Merck Millipore)–water. Buffer A was 0.1% formic acid, and buffer B was 80% acetonitrile□plus□0.1% formic acid. The HPLC pumped a flow-rate of 400 nL/min with a programmed gradient from 5-34% solvent B (A□=□0.1% formic acid; B□=□80% acetonitrile, 0.1% formic acid) over 45 min, followed by a gradient from 34% B to to 90% B in 2 min and 5 min isocratic at 90% B. The HPLC system was coupled online with a Q Exactive HF mass spectrometer (Thermo Fisher Scientific, San Jose, CA) acquiring data in a data-independent acquisition (DIA) mode as previously optimized^77,78^. Specifically, DIA consisted of a full scan MS spectrum (m/z 300−1100) followed by 16 MS/MS with windows of 50□m/z and detected all in high resolution. The full scan MS was acquired with a resolution of 60,000 and an automatic gain control (AGC) target of 1×10^6^. MS/MS was performed with and AGC target of 5×10^5^ using higher-collision dissociation with normalized collision energy of 27.

DIA data obtained from the bottom-up analysis were searched using EpiProfile 2.0^79^ and validated in Skyline (MacCoss Lab). The peptide relative ratio was calculated using the total area under the extracted ion chromatograms of all peptides with the same amino acid sequence (including all of its modified forms) as 100%. For isobaric peptides, the relative ratio of two isobaric forms was estimated by averaging the ratio for each fragment ion with different mass between the two species.

### Analysis of ACSS2i bioavailability

Mice (n=2 for each timepoint) were intraperitoneally injected with cAcss2i at a dosage of 0.5 mg/kg, and sacrificed 10 minutes, 20 minutes and 30 minutes after administration. Brains were rapidly collected and snap-frozen with liquid nitrogen. Hippocampi were free-hand dissected on ice, and bilaterally pooled. A small amount of LC/MS-grade water was added to bring up the total amount of water in the sample to ∼80%. Samples were homogenized by mechanical disruption in a 1:2 ratio of chloroform and methanol. Additional chloroform was to bring the chloroform:methanol ratio to 1:1, and the sample was vigorously vortexed. Finally, water was added to the homogenate such that the final concentration of water:chloroform:methanol ratio was equivalent (with the initial water content of the hippocampal sample counting towards the total water volume). Samples were once again vigorously vortexed, then passed through a 0.22 micron filter. To separate the aqueous and organic layers, samples were centrifuged for 5 minutes at 10000 rpm in a tabletop microcentrifuge. The lighter aqueous layer was carefully removed, and the lower organic phase (containing the hydrophobic ACSS2i) was dried using a ThermoSavant SC110A SpeedVac Plus (Thermo Scientific). Once ready to inject in the MS, samples were reconstituted in Buffer A (0.1% formic acid). Rat pharmacokinetic data for cACSS2i and nACSS2i was similarly generated by Charles River Laboratories. Briefly, cACSS2i and nACSS2i were administered intravenously, and plasma and brains were collected at the indicated timepoints. Concentrations of both compounds remaining at each timepoint were determined by LC/MS/MS.

### MS analysis of ACSS2i

Samples were injected onto 75□μm ID × 25□cm Reprosil-Pur C18-AQ (3□μm; Dr. Maisch GmbH, Germany) nano-column packed in-house using an EASY-nLC nanoHPLC (Thermo Fisher Scientific). The nanoLC pumped a flow-rate of 400 nL/min with a programmed gradient from 0% to 70% solvent B (A□=□0.1% formic acid; B□=□80% acetonitrile, 0.1% formic acid) over 5□minutes, followed by a gradient from 70% to 100% solvent B in 10□minutes and 5□min isocratic at 80% B. The instrument was coupled online with a Q-Exactive (Thermo Fisher Scientific) mass spectrometer acquiring data in a Parallel Reaction Monitoring (PRM) mode. Specifically, this method consisted of a full scan MS spectrum (350-450 m/z) followed by a PRM scan targeting 411.09490 m/z, which corresponds to the cACSS2i. The full scan MS was acquired with a resolution of 70,000 and an automatic gain control (AGC) target of 3×10^6^. The PRM scan was performed with and AGC target of 2×10^5^ using higher-collision dissociation with normalized collision energy of 27. The PRM data obtained was analyzed manually, where the peak areas corresponding to the cACSS2i were extracted.

## ACKNOWLEDGEMENTS

We thank E. Korb and K. Wellen for insightful discussions on behavior and metabolism, respectively. Acss2^KO^ were generated with significant help from the University of Pennsylvania Transgenic and Chimeric Mouse Core (IDOM, the Center for Molecular Studies in Digestive and Liver Diseases). Behavior procedures were performed with assistance from The Neurobehavior Testing Core at UPenn/ITMAT and IDDRC at CHOP/Penn, U54 HD086984. We thank J. A. Youssefian for material and logistical support. This work was supported by NIH RO1AA027202 (S.L.B), T32 GM-07229 (D.C.A), F31 CA247348-02 (M.M.), AARF-19-618159 (G.E.),

## Notes

### Competing Interest Statement

S.L.B. and P.M. are co-founders of EpiVario, Inc. EpiVario provided experimental compounds (ACSS2i) through a sponsored research agreement with the University of Pennsylvania.

