## Supplemental Figures for "Targeting acetyl-CoA metabolism attenuates the formation of fear memories through reduced activity-dependent histone acetylation"

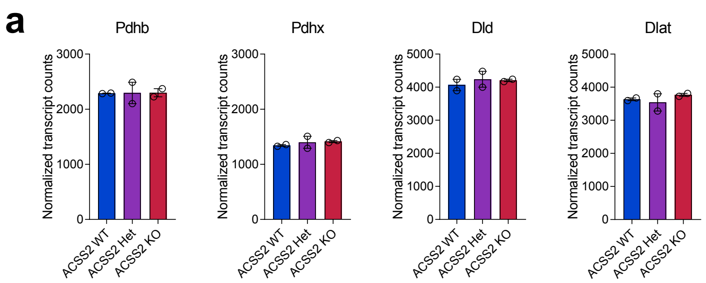


**Supplementary Figure 1:** Acss2KO mice show no change in the expression of PDC subunits. **a)** Bar charts showing mRNA expression levels of PDC subunits (Pdhb, Pdhx, Dld, Dlat). mRNA levels shown as normalized transcript counts.

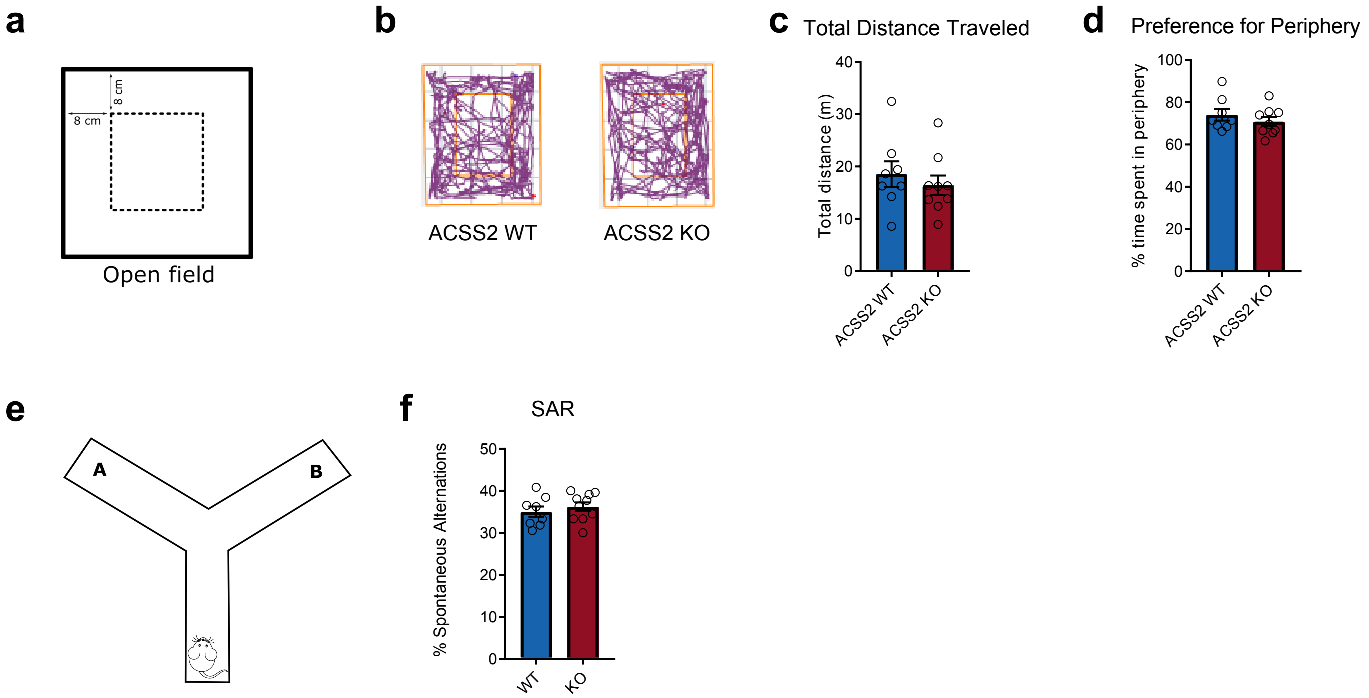


**Supplemental figure 2:** ACSS2 KO mice perform normally under baseline conditions. **a)** Open field (OF) assay diagram. **b)** Representative paths during OF for WT and Acss2 KO mice visualized using AnyMaze motion tracking software. **c)** OF locomotion quantified by total distance in meters (WT: n=8; Acss2 KO: n9; p = n.s. (0.5777), unpaired t-test). **d)** OF thigmotaxis quantified by time spent in periphery over total time (WT: n=8; Acss2 KO: n9; p = n.s. (p = 0.6315), student's unpaired t-test). **e)** Y-maze test of spontaneous alternation diagram. **f)** Percent spontaneous alternations in y-maze described as number of successful spontaneous alternations over total arm entries (WT: n=8; Acss2 KO: n=9; p = n.s, unpaired t-test)

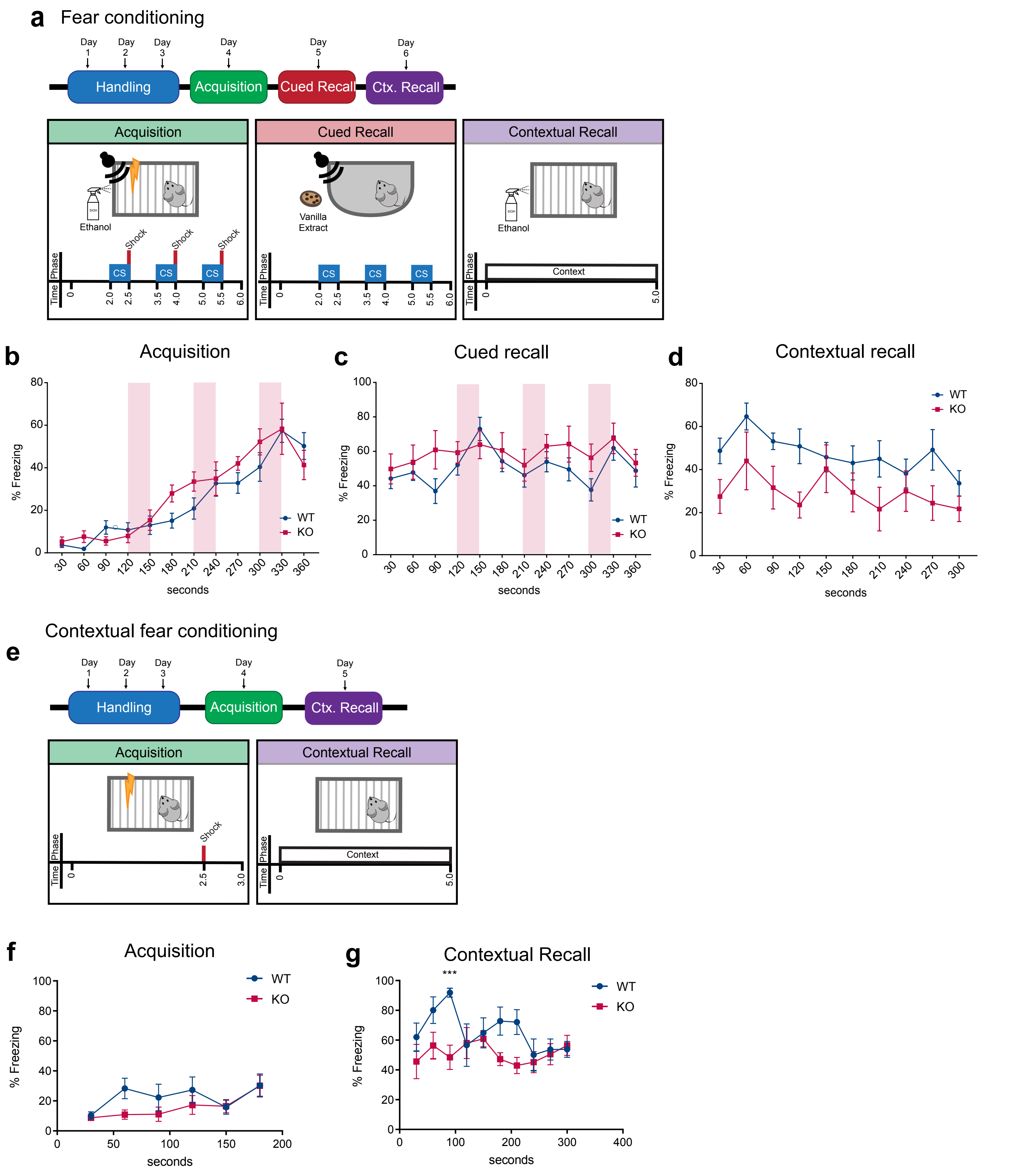


**Supplemental Figure 3:** Acss2$^{KO}$ mice exhibit reduced fear conditioning. **a)** Fear conditioning schematic, showing training/ acquisition (panel 1),cued/auditory recall (panel 2), and contextual recall (panel 3). **b)** Fear response during training/ acquisition reflected as % time freezing, binned into 30 s intervals. Tone-on indicated by transparent red bars; shocks, by black arrows. (p = n.s. between WT and Acss2 KO, 2-way ANOVA). **c)** Fear response during cued/auditory recall reflected as % time freezing, binned into 30 s intervals. Tone-on indicated by transparent red bars; no shock. **d)** Fear response during contextual recall reflected as % time freezing/total time, binned into 30 s intervals. (p = 0.0305, 2-way ANOVA)**. e)** Contextual conditioning schematic, showing training/acquisition (panel 1), and contextual recall (panel 2). **f)** Fear response during training/acquisition reflected as % time freezing/total time, binned into 30s intervals. **g)** Fear response during contextual recall reflected as % time freezing/total time, binned into 30s intervals (p=0.0057, 2-way ANOVA)

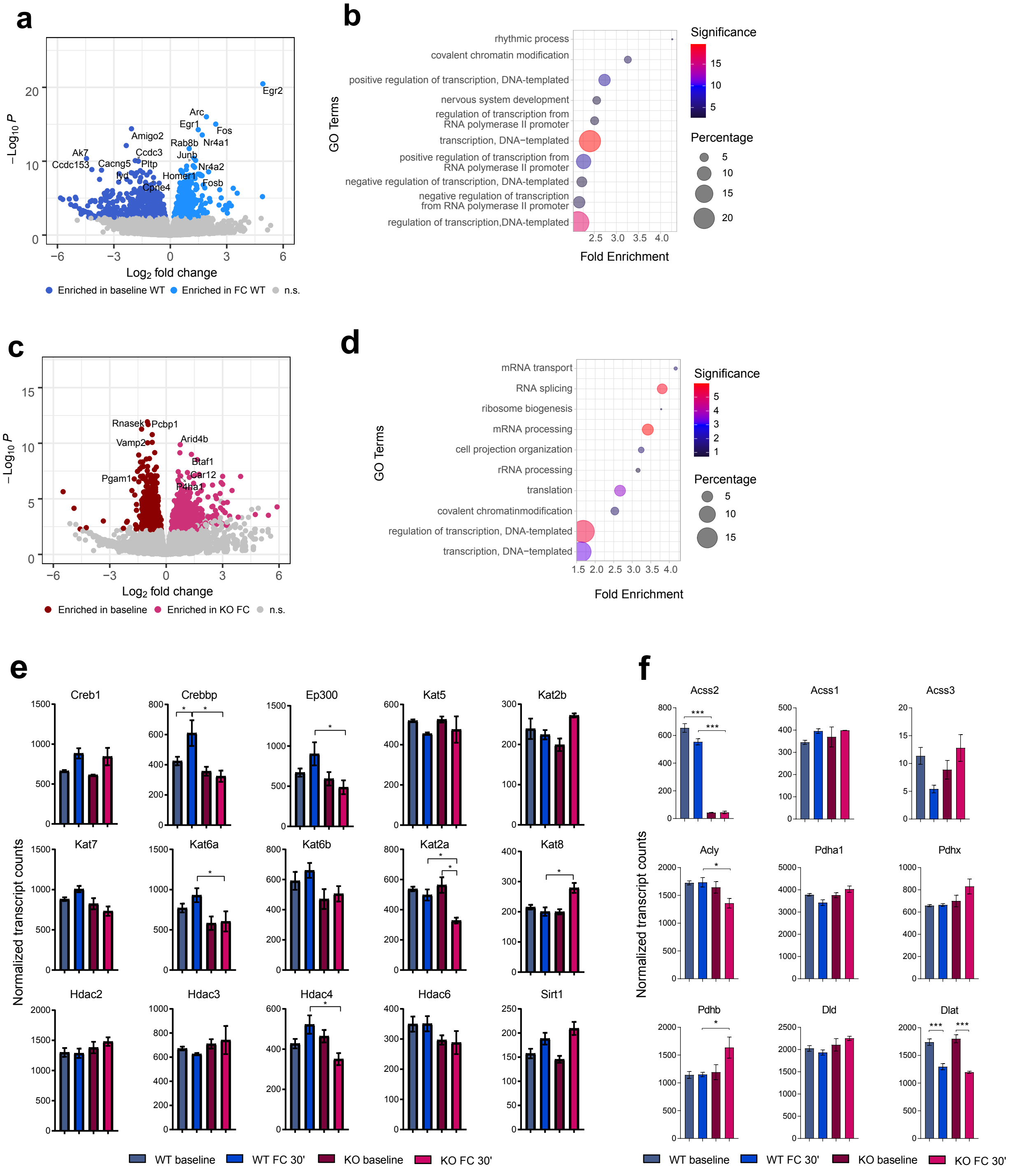


**Supplemental Figure 4:** Loss of ACSS2 negatively impacts the transcription of activity-dependent genes required for LTM formation in a fear conditioning model

**a)** Volcano plot for gene expression changes in WT mice 30 minutes after fear conditioning. (WT baseline: n=5; WT FC 30': n = 3; padj < 0.05)

**b)** Top 10 GO terms (DAVID, BP) enriched in WT FC 30' (DAVID, BP) compared to WT baseline.

**c)** Volcano plot for gene expression changes in KO mice 30 minutes after fear conditioning. (KO baseline: n=3; KO FC 30': n = 2; padj < 0.05)

**d)** Top 10 GO terms (DAVID, BP) enriched in KO FC 30' (DAVID, BP) compared to KO baseline.

**e)** Bar plots of histone acetyltransferases and histone deacetylases showing normalized transcript counts across all four conditions (RNA-seq; Deseq2).

**f)** Bar plots for acetyl-CoA-producing metabolic enzymes showing normalized transcript counts across all four conditions (RNA-seq; Deseq2).

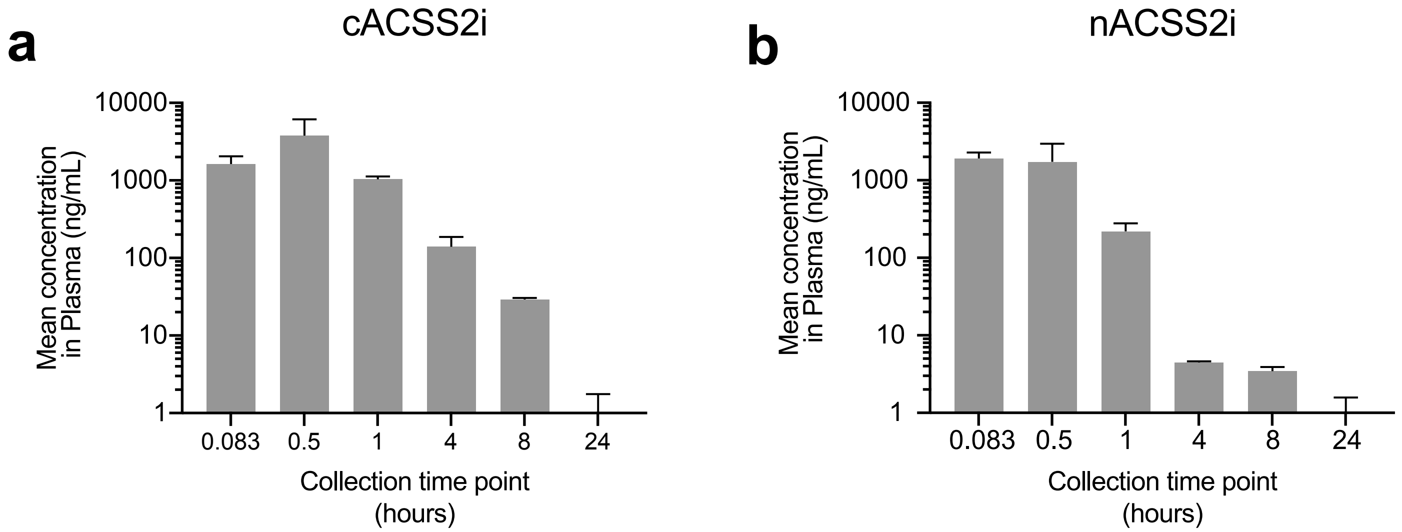


**Supplemental Figure 5:**
Pharmacokinetic assay in rats for cACSS2i and nACSS2i in plasma following IV administration.
a) cACSS2i is detected in blood plasma of rats at time points after IV administration. cACSS2i is cleared from the animal by the 24 hour mark.
b) nACSS2i is detected in blood plasma of rats at time points after IV administration. Relative to cACSS2i, the nACSS2i is detected at lower amounts at the 4 hours and 8 hours time points.

a

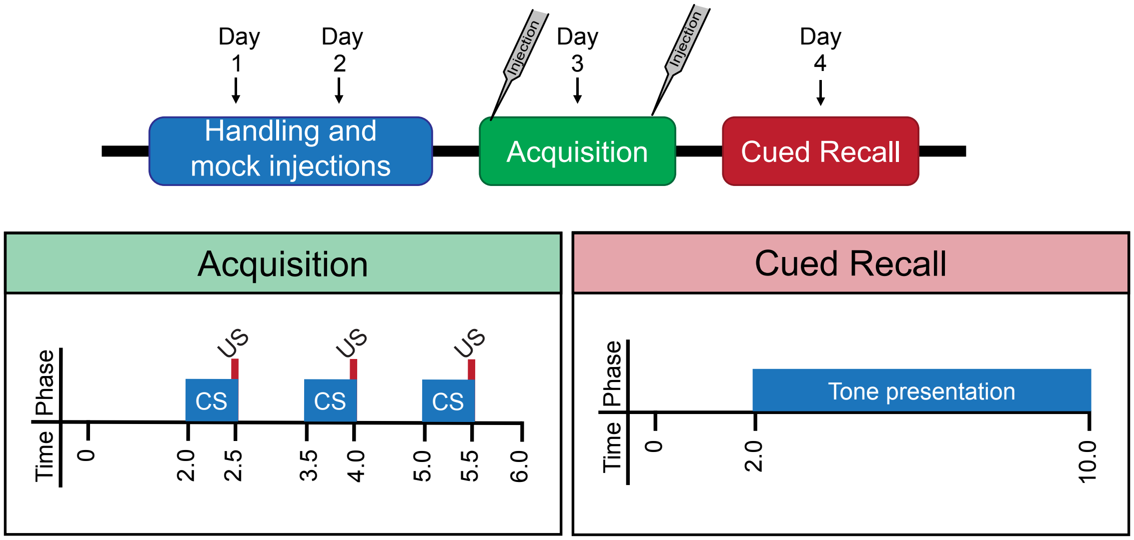


b

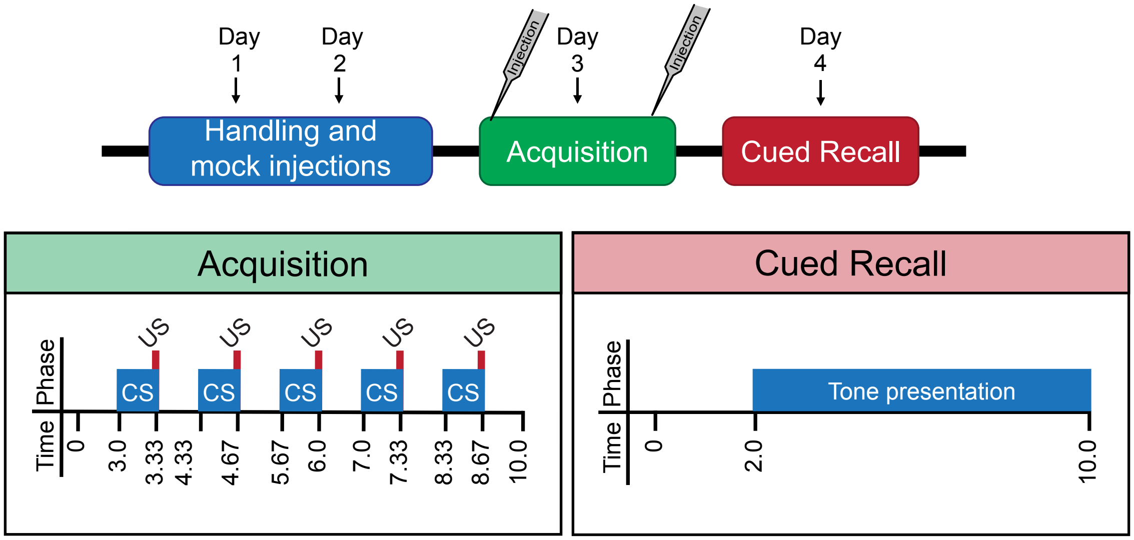


**Supplemental Figure 6:**
Schematics outlining cued fear conditioning paradigm used in mouse and rat.
a) In mouse, we utilized a 3 conditioned stimululs-unconditioned stimulus pairing model at acquisition. At cued recall, mice were exposed to a prolonged 8min tone.
b) Rats were exposed to 5 pairings of the conditioned stimululs-unconditioned stimulus at acquisition. At cued recall, rats were exposed to a prolonged 8min tone.

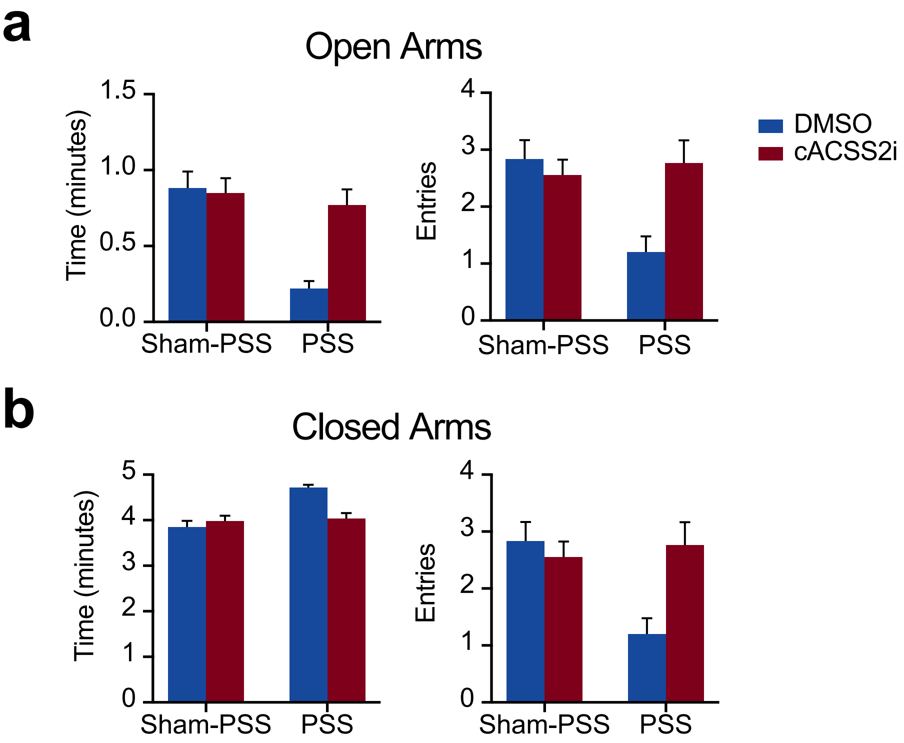


**Supplemental Figure 7:**
Measurements of activity in elevated plus maze following PSS exposure in rats.

a) Time spent in open arms of EPM is significantly reduced in PSS-exposed animals treated with DMSO. This indicates an increase in anxiety. In contrast, cACSS2i-treated PSS-exposed rats spend similar amounts of time in open arms as Sham-PSS animals. This trend is similarly observed when counting entries into open arms.

b) Time spent in closed arms of EPM is increased in PSS-exposed animals treated with DMSO. In contrast, cACSS2i-treated PSS-exposed rats spend similar amounts of time in closed arms as Sham-PSS animals. This trend is similarly observed when counting entries into closed arms.
